# Common substructures and sequence characteristics of sandwich-like proteins from 42 different folds

**DOI:** 10.1101/2020.05.27.108969

**Authors:** A.E. Kister

## Abstract

This study addresses the following fundamental question: Do sequences of protein domains with sandwich architecture have common sequence characteristics even though they belong to different superfamilies and folds? The analysis was carried out in two stages: determination of substructures in the domains that are common to all sandwich proteins; and detection of common sequence characteristics within the substructures. Analysis of supersecondary structures in domains of proteins revealed two types of four-strand substructures that are common to sandwich proteins. At least one of these common substructures was found in proteins of 42 sandwich-like folds (as per structural classification in the CATH database). Comparison of the sequence fragments corresponding to strands that make up the common substructures revealed specific rules of distribution of hydrophobic residues within these strands. These rules can be conceptualized as grammatical rules of beta protein linguistics. Understanding of the structural and sequence commonalities of sandwich proteins may also be useful for rational protein design.

## Introduction

According to the protein structural hierarchical classification adopted in the CATH database, ‘architecture level’ is a general description of the gross arrangement of secondary structures that is independent of their connectivity.^1^ Thus, a particular architecture may describe structures from non-homologous superfamilies with different numbers of secondary structure elements and diverse connectivity. Examples of protein architecture are beta-sandwich or beta-barrel. An important unanswered question is whether any specific sequence features are responsible for the specific protein architecture. This study focuses on the relationship between sequences and protein structures with sandwich-like architecture. Sandwich folds in beta-protein domains are characterized by two main beta sheets packed against each other. This architecture characterizes more than 40 protein folds and 532 superfamilies in the CATH database structural classification of protein domains. Immunoglobulin-like and Jelly Rolls folds have the largest number of superfamilies: 318 and 127, respectively.

Usual methods for detecting sequence similarity using similarity scores ^2–4^ have not so far been successful in uncovering common sequence characteristics in the highly non-homologous sandwich proteins. In this study, an alternative, two-stage approach to uncovering sequence similarities is proposed. The first stage is to identify ‘substructures’ common to proteins of all sandwich folds. Substructures are defined as specific arrangements of two pairs of consecutive strands in two main beta-sheets. The four-strand substructure represents the smallest possible sandwich structure. One example of such a substructure – ‘the interlock’ - was previously shown to be present across a group of sandwich-beta proteins from different superfamilies and protein folds.^5^ Here we describe another type of conserved arrangement of four strands within sandwich-like folds proteins. We show that at least one of these two types of substructure is always present in sandwich-like proteins. Since these substructures are an invariant feature of sandwich proteins, their formation is likely a necessary condition for the development of structures with this architecture.

The second stage of analysis is to find common sequence characteristics within the strands that make up the conserved substructures (the ‘substructure strands’). Numerous lines of evidence point to the decisive role of hydrophobic residues in beta-sheet formation of beta structures. Hydrophobic residues (a) have a special propensity for beta sheets;^6–8^ b) play a key role in the formation of ‘hydrophobic zipper hypothesis’;^9^ c) are mostly responsible for protein folding and stability;^10–11^ and d) the development of hydrophobic core;^12–14^; and, e), are also important for the formation of β-sheet packing interfaces.^15^ Therefore, the analysis of substructure strands focuses on the locations and contacts of hydrophobic residues in sequences. Because contacts between oppositely charged residues – ‘salt bridges’ - also significantly contribute to protein stability,^16–17^ these interactions were also taken into account in our analysis.

As the standard alignment procedures did not reveal conserved hydrophobic positions in sequences from different superfamilies, our analysis focused on the finding of pairs of hydrophobic residues that form contacts between substructure strands. We identified such contacts that are invariably present in substructures of sandwich-like proteins. The matrix of conserved contacts can be regarded as the common sequence characteristic of sandwich proteins. Another characteristic feature of the substructures is the relatively large proportion of hydrophobic residues in the substructure strands.

## Analysis and Results

### Two types of conserved supersecondary substructures in sandwich domains

<u>Substructure type 1</u> is formed by four consecutive strands. Two strands (*i*-1 and *i*) form backbone hydrogen bond contacts in one sheet and the other two strands (*i*+1 and *i*+2) form hydrogen bond contacts in the other sheet (Fig. 1). Two variants of this substructure, subtype 1a, and 1b, differ in the relative locations of strands *i*-1 and *i*+2 as shown in Fig. 1a and 1b, respectively. Substructure 1a was found in 20 folds and substructure 1b - in 10 folds (Table 1, columns “SubS 1-4”).

**Figure 1.**
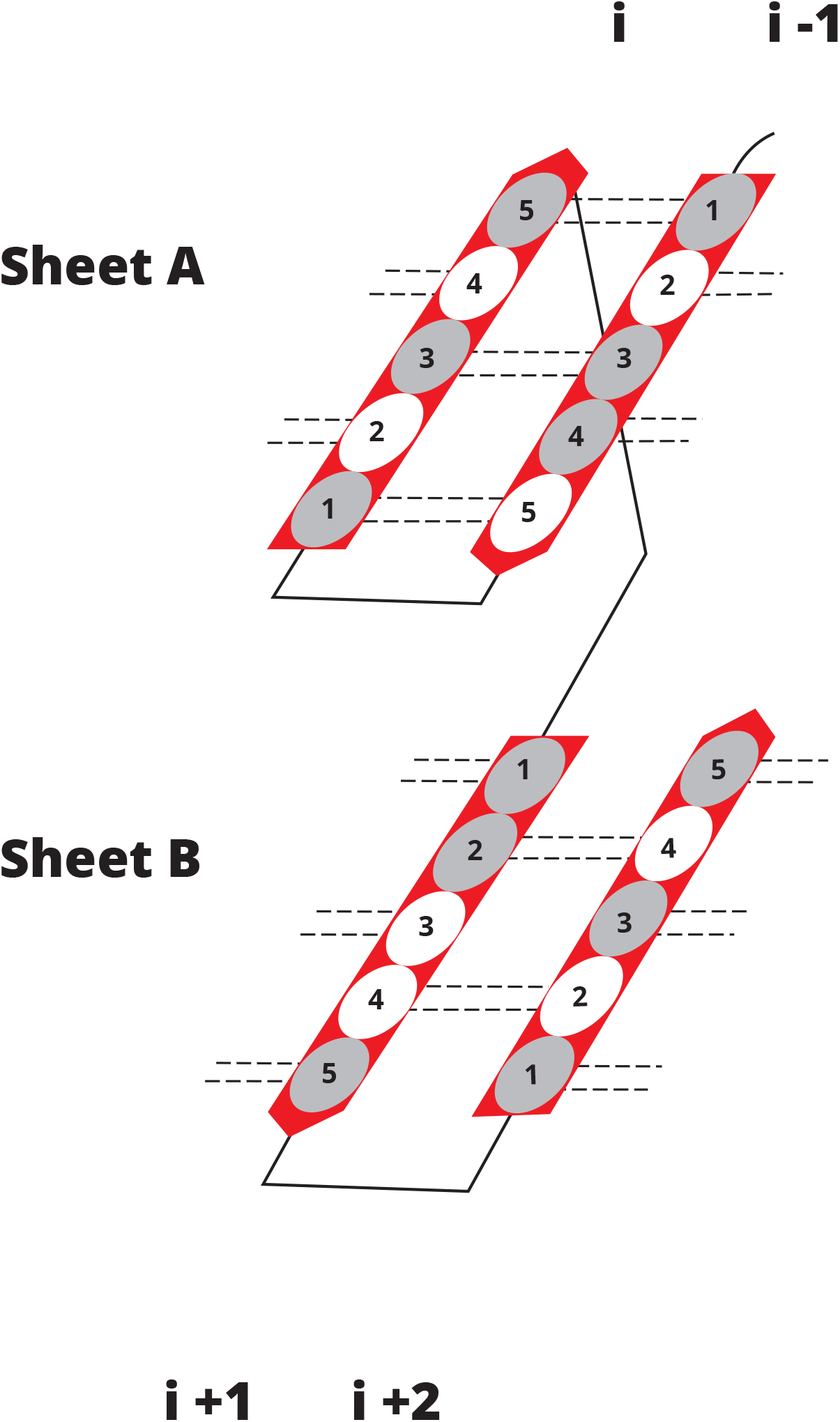
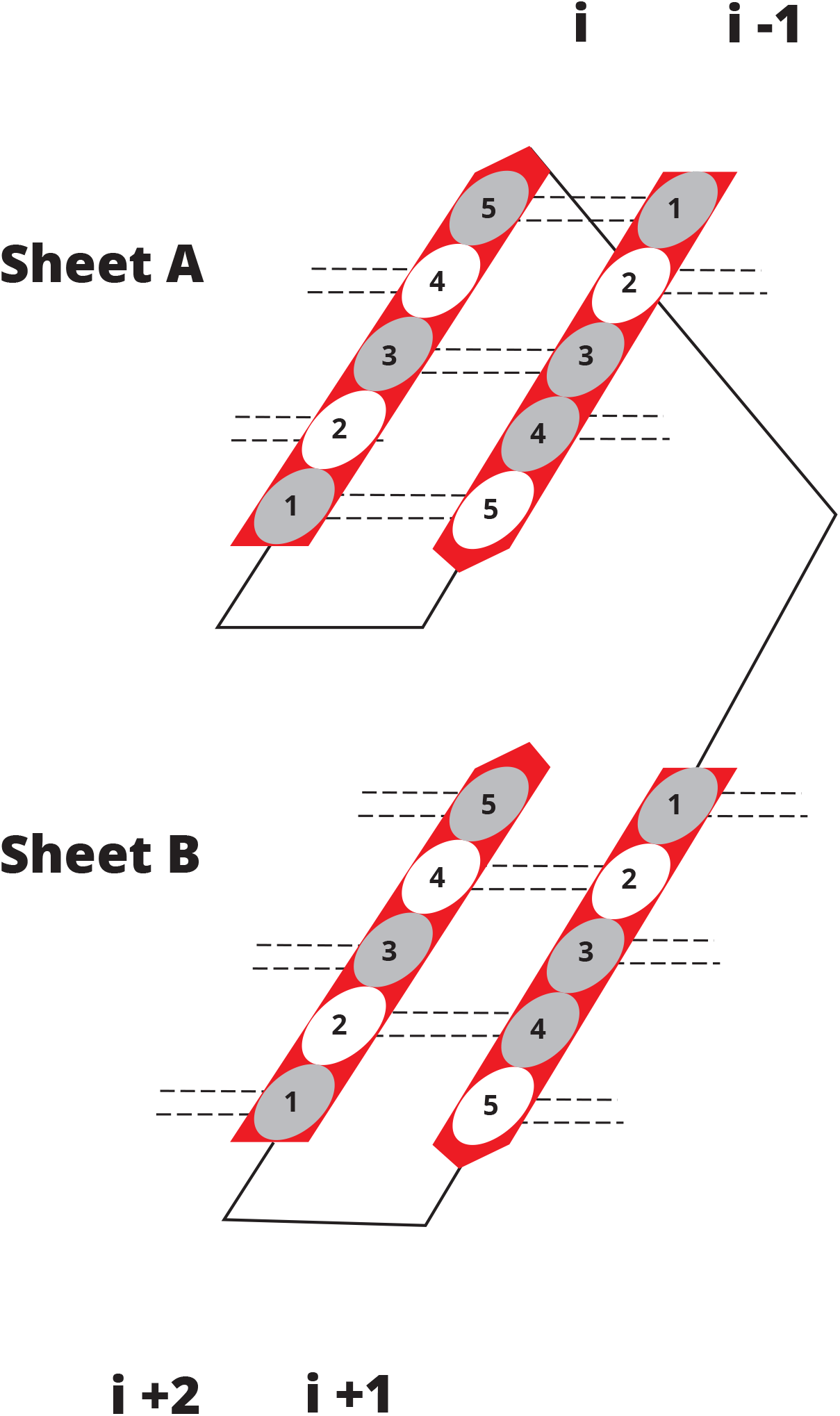
Conserved Substructure type 1. Two variants of Substructures with specific arrangements of strands *i*-1, *i*, *i*+1, and *i*+2 (1a and 1b) are shown. Beta strands are represented by arrows and protein loops by lines. Ovals in the arrows represent residues in strands. Five positions of residues in each strand are considered. In this model, it is assumed that residues at positions 1, 3, and 5 are directed inside, while residues at positions 2 and 4 are directed outward. The ‘grey positions’ are occupied by hydrophobic residues.

**Table 1:** Sandwich domains in 42 different folds: Supersecondary structures. Substructures. Columns: **CATH folds**: Classification of protein folds according to CATH database; Sfam: Numbers of superfamilies in the given fold; chain forming a sandwich-like domain**; SSS domain:** Supersecondary structure of a sandwich like domain. The order in SSS PDB code: PDB code of the structure that represents the protein structures in this fold; **Chain fragment:** Fragment of the polypeptide **chain forming** corresponds to the arrangement of the strands in structures. “=” - backbone hydrogen bond contacts between strands in a beta-sheet. Two comments for SSS are presented if 1) two consecutive strands x and x + 1 are located in different beta-sheets and are not described by any substructure (SubS) {see, for example, structure 1occ, CATH fold 2.60.11: there are no substructure with two strands #1 and 2 in this domain} and 2) two neighboring strands in a beta-sheet are parallel {see, for example, structure 1occ the strands #1 and 5}. In the columns **SubS 1-4** are shown 1) the classification of all substructures in the domain; 2) two pairs of strands in sheets A and B that form the substructure; 3) the total number of PHRs in the substructure + if exist, the number of pairs of oppositely charged residues (for example, in structure 4f01, CATH folds 2.60.34 in SubS 1b PHR is 6 + 2 pairs of oppositely charged residues; 4) the total number of hydrophobic residues in the substructure in round brackets (in the same SubS the number is equal to 15); 5) the total number of PCRs in the substructure in square brackets [in the same substructure the number is 15]. * In this paper, the cyclic numbering of strands in SSS is adopted: the next strand after the last strands in SSS is strand #1. Thus the last and first strands in sandwich domains are considered to be consecutive. See, for example, substructure 2c in structure 1g6e CATH folds 2.60.30.

| CATH folds | Sfam | PDB code | Chain fragment | SSS domain | SubS 1 | SubS 2 | SubS 3 | SubS 4 |
| --- | --- | --- | --- | --- | --- | --- | --- | --- |
| 2.60.9 | 1 | 2hnu | A: 7-87 | sheet A: 4=3=2=1<br>sheet B: 5=6=7=8 | 1a<br>4=3<br>5=6<br>3(8)[8] | 1a *<br>7=8<br>2=1<br>2 (7)[8] | — | — |
| 2.60.11 | 1 | 1occ | F:1-98 | sheet A: 4=5=1<br>sheet B: 3=2<br>No SubS for strands # 1 and 2<br>strands # 1 and 5 are parallel | 1a<br>4=5<br>3=2<br>4(14)[13] | — | — | — |
| 2.60.15 | 1 | 1h8e | H: 15-103 | sheet A: 2=1=9=8=5<br>sheet B: 4=3=6=7=10<br>strands #1 and 9 parallel<br>No SubS for strands # 4 and 5 | 1b<br>2=1<br>4=3<br>4 (13)[14] | 1a<br>9=8<br>6=7<br>4 (14) [14] | 2c<br>8=5<br>6=7<br>2 (14)[14] | 2c<br>9=8<br>7=10<br>4 (12)[11] |
| 2.60.20 | 4 | 4iau | A: 80-162 | sheet A: 2=1=3=6<br>sheet B: 7=4=5 | 2d*<br>1=3<br>7=4<br>3+2 (10)[11] | 2a<br>3=6<br>7=4<br>3+2(9)[9] | 2d<br>3=6<br>4=5<br>2 (8) [8] | — |
| 2.60.30 | 1 | 1g6e | A:1-87 | sheet A: 1=3=2=5=8<br>sheet B: 4=9=6=7 | 2c*<br>1=3<br>4=9<br>1 (7)[8] | 2d<br>5=8<br>4=9<br>2 (7) [8] | 2a<br>5=8<br>9=6<br>5(12)[14] | 2d<br>5=8<br>6=7<br>3(11) [13] |
| 2.60.34 | 3 | 4f01 | B:382-506 | sheet A: 5=4=1=2<br>sheet B: 8=7=6=3<br>No SubS for strands # 2 and 3 | 1b<br>5=4<br>7=6<br>6+2(15) [15] | 2d<br>5=4<br>6=3<br>4+1(13)[15] | — | — |
| 2.60.40. | 318 | 1dr9 | A:1-105 | sheet A: 1=9=8=3=4=5<br>sheet B: 2=7=6<br>No SubS for strands # 1 and 2 | 1a<br>4=5<br>7=6<br>6 (15)[10] | 2a<br>8=3<br>2=7<br>7(18)[15] | — | — |
| 2.60.60 | 5 | 1rrh | A:8-167 | **sheet A: 4=5=1=7=8**<br>**sheet B: 3=2=6=9** | **1a**<br>**4=5**<br>**3=2**<br>**4 (14)[12]** | **2a**<br>**5=1**<br>**2=6**<br>**6+1(19)[18]** | **2d**<br>**7=8**<br>**6=9**<br>**6+1(15)[13]** | — |
| 2.60.90 | 5 | 1knb | A:182-365 | **sheet A: 10=3=2=1=5=6**<br>**sheet B: 7=8=9=4**<br>strands #3 and 4 are parallel<br>No SubS for strands # 4 and 5 | **2c**<br>**10=3**<br>**9=4**<br>**6(17)[14]** | — | — | — |
| 2.60.98. | 6 | 1svb | A:1-51,<br>135-192 | **sheet A: 1=3=4=5=6**<br>**sheet B: 2=9=8=7**<br>No SubS for strands # 1 and 2<br>strands #1 and 3 are parallel | **1a**<br>**5=6**<br>**8=7**<br>**3 +1 (13)[13]** | **2a\***<br>**1=3**<br>**2=9**<br>**4 + 1 (14) [14]** | — | — |
| 2.60.110 | 1 | 2vhk | A:1-207 | **sheet A: 4=1=11=8=9**<br>**sheet B: 3=2=5=6=7=10** | **2d**<br>**4=1**<br>**3=2**<br>**4(13)[13]** | **2a**<br>**4=1**<br>**2=5**<br>**6(14)[14]** | **2a**<br>**11=8**<br>**7=10**<br>**2(6)[7]** | **2d**<br>**8=9**<br>**7=10**<br>**3(8)[10]** |
| 2.60.120 | 127 | 3HIF | A:7-130 | **sheet A: 1=8=3=6**<br>**sheet B: 2=7=4=5** | **2 d**<br>**1=8**<br>**2=7**<br>**3+2(8)[7]** | **2a**<br>**8=3**<br>**2=7**<br>**5(16)[12]** | **2a**<br>**3=6**<br>**7=4**<br>**6(15)[10]** | **2d**<br>**3=6**<br>**4=5**<br>**7(17)[10]** |
| 2.60.130 | 7 | 1dlm | A:3-311 | **sheet A: 1=5=4=7=8=9**<br>**sheet B: 2=6=3=11=10** | **2b**<br>**1=5**<br>**2=6**<br>**2 (9)[10]** | **2d**<br>**5=4**<br>**6=3**<br>**2(14)[15]** | **2a**<br>**4=7**<br>**6=3**<br>**3(16)[16]** | **1a**<br>**8=9**<br>**11=10**<br>**3(12)[9]** |
| 2.60.169 | 1 | 2bpa | 1: 1-420 | **sheet A :1=8=3=6**<br>**sheet B: 2=7=4=5** | **2d**<br>**1=8**<br>**2=7**<br>**3(22)[23]** | **2a**<br>**8=3**<br>**2=7**<br>**4+1(23)[24]** | **2d**<br>**3=6**<br>**4=5**<br>**7+1(26)[24]** | **2a**<br>**3=6**<br>**7=4**<br>**5+1(25)[24]** |
| 2.60.175 | 2 | 4mby | H:25-300 | **sheet A: 1=9=3=6**<br>**sheet B: 2=7=4=5**<br>**sheet B: 8=7**<br>No SubS for strands # 1 and 2 | **2d**<br>**3=6**<br>**4=5**<br>**5(15)[16]** | **2a**<br>**3=6**<br>**7=4**<br>**6(18)[16]** | **1b\***<br>**1=9**<br>**8=7**<br>**4(13)[14]** | **2d**<br>**3=6**<br>**2=7**<br>**5(16)[14]** |
| 2.60.200. | 6 | 4r9p | A:27-236 | **sheet A: 2=1=10=9=7**<br>**sheet B: 4=3=5=6=8**<br>strands #3 and 4 are parallel | **1a**<br>**2=1**<br>**4=3**<br>**6 (17)[13]** | **2a**<br>**9=7**<br>**6=8**<br>**5 (15)[12]** | — | — |
| 2.60.210 | 1 | 1lb4 | A:348-505 | **sheet A: 1=8=5=6**<br>**sheet B: 2=3=4=7**<br>No SubS for strands # 1 and 2 | **1b**<br>**5=6**<br>**3=4**<br>**4 (19)[18]** | **2d**<br>**5=6**<br>**4=7**<br>**4 (17)[16]** | **2a**<br>**8=5**<br>**4=7**<br>**7 + 1 (18) [16]** | — |
| 2.60.220 | 6 | 1hn0 | A:900-1020 | **sheet A: 1=2=3=6=7=12=9**<br>**sheet B: 4=5=8=11=10** | **2d**<br>**3=6**<br>**4=5**<br>**4(11)[12]** | **2d**<br>**6=7**<br>**5=8**<br>**3(13)[13]** | **2a**<br>**12=9**<br>**8=11**<br>**2(9)[14]** | **2d**<br>**12=9**<br>**11=10**<br>**2(8)[13]** |
| 2.60.240 | 3 | 1cq3 | A:1-224 | **sheet A: 10=9=1=6=5**<br>**sheet B: 8=7=4=3=2**<br>strands # 7 and 8 are parallel<br>No SubS for strands # 1 and 2 | **1b**<br>**10=9**<br>**8=7**<br>**5+3(17)[19]** | **2d**<br>**6=5**<br>**7=4**<br>**5(14)[16]** | — | — |
| 2.60.250 | 1 | 1p35 | A:2-300 | **sheet A: 1=4=7=6**<br>**sheet B: 10=11=12=13=9=2=3=8=5** | **2d**<br>**1=4**<br>**2=3=**<br>**7(16)[17]** | **2a**<br>**4=7**<br>**8=5**<br>**8+1(18)[13]** | **2d**<br>**7=6**<br>**8=5**<br>**5+1(15)[13]** | — |
| 2.60.260 | 4 | 3tj8 | A:1-75 | **sheet A: 1=5=4**<br>**sheet B: 2=6=7=3**<br>No SubS for strands # 3 and 4<br>strands #1 and 5 are parallel<br>strands #3 and 7 are parallel | **2b**<br>**1=5**<br>**2=6**<br>**4+1(11) [9]** | **1a**<br>**5=4**<br>**6=7**<br>**5+1(15)[14]** | — | — |
| 2.60.270. | 4 | 1gwy | A:1-175 | **sheet A: 1=5=2=12=11=10**<br>**sheet B: 4=3=6=7=8=9**<br>strands #2 and 12 are parallel | **2d**<br>**5=2**<br>**4=3**<br>**4+1(12)[12]** | **2a**<br>**5=2**<br>**3=6**<br>**8(23)[18]** | **1a**<br>**11=10**<br>**8=9**<br>**3+1(9)[11]** | — |
| 2.60.290 | 2 | 1nc7 | A:0-115 | **sheet A: 9=1=6=3=4**<br>**sheet B: 8=7=2=5** | **1a\***<br>**9=1\***<br>**8=7**<br>**4+2(13)[14]** | **2a**<br>**1=6**<br>**7=2**<br>**6+1(17)[17]** | **2d**<br>**1=6**<br>**2=5**<br>**7+2(20)[18]** | **2d**<br>**3=4**<br>**2=5**<br>**5+2(15)[15]** |
| 2.60.300 | 1 | 2apn | A: 9-104 | **sheet A: 1=6=7**<br>**sheet B: 3=2=5=4**<br>strands # 1 and 6 are parallel<br>strands # 2 and 5 are parallel | **2c**<br>**1=6**<br>**2=5**<br>**4 (11) [11]** | **1a**<br>**6=7**<br>**5=4**<br>**4 (10) [11]** | — | — |
| 2.60.310 | 2 | 1js8 | B:2787-2892 | **sheet A: 7=1=4**<br>**sheet B: 6=5=2=3** | **1a**<br>**7=1**<br>**6=5**<br>**4 (13) [11]** | **2a**<br>**1=4**<br>**5=2**<br>**7+2 (19) [16]** | **2d**<br>**1=4**<br>**2=3**<br>**7 (17) [14]** | — |
| 2.60.320 | 3 | 3ld7 | A:39-125 | **sheet A: 2=1=7=6**<br>**sheet B: 3=4=5** | **1a**<br>**2=1**<br>**3=4**<br>**4 (15) [15]** | **1a**<br>**7=6**<br>**4=5**<br>**4+2 (14) [15]** | — | — |
| 2.60.330 | 1 | 1n7v | A: 43-66<br>A: 321-419 | **sheet A: 1=6=7=8=9**<br>**sheet B: 5=2=3=4**<br>strands # 1, 2, 6 are parallel | **2c**<br>**1=6**<br>**5=2**<br>**3 (9) [9]** | — | — | — |
| 2.60.340 | 1 | 1n80 | A:12-87,<br>246-334 | **sheet A : 7=5=1=2=6**<br>**sheet B: 8=4=3** | **1a**<br>**1=2**<br>**4=3**<br>**4(14)[12]** | **2d**<br>**7=5**<br>**8=4**<br>**5(15)[16]** | — | — |
| 2.60.350 | 1 | 1x0c | A: 18-189 | **sheet A: 4=5**<br>**sheet A: 1=5=11=10=7**<br>**sheet B: 3=2=6=9=8** | **2b**<br>**1=5**<br>**2=6**<br>**4(12)[14]** | **1a**<br>**4=5**<br>**3=2**<br>**3+1(8)[8]** | **2a**<br>**10=7**<br>**6=9**<br>**5(12)[13]** | **2d**<br>**10=7**<br>**9=8**<br>**5(12)[11]** |
| 2.60.370 | 1 | 1so9 | A:21-151 | **sheet A: 3=1=7=5**<br>**sheet B: 2=4=6**<br>No SubS for strands # 2 and 3 | **2a**<br>**3=1**<br>**2=4**<br>**4+2(15)[15]** | **2a**<br>**7=5**<br>**4=6**<br>**7+2(19)[17]** | — | — |
| 2.60.390 | 1 | 1tg7 | A: 577-665 | **sheet A: 1=4=8=7**<br>**sheet B: 2=3=5=6**<br>No SubS for strands # 4 and 5 | **2d**<br>**1=4**<br>**2=3**<br>**6+1(14)[13]** | **1a**<br>**8=7**<br>**5=6**<br>**1(5)[6]** | — | — |
| 2.60.410 | 1 | 1hd8 | A:263-356 | **sheet A: 4=1**<br>**sheet B: 5=7=8=3=2=6**<br>No SubS for strands # 4 and 5 | **2d**<br>**4=1**<br>**3=2**<br>**3(15)[15]** | **2a**<br>**4=1**<br>**8=3**<br>**4(14)[12]** | — | — |
| 2.60.420 | 1 | 1h54 | A:310-315,<br>685-753 | **sheet A: 1=4=5=6**<br>**sheet B: 2=3=7=8** | **2d**<br>**1=4**<br>**2=3**<br>**4(11)[12]** | **1b**<br>**5=6**<br>**7=8**<br>**5+2(15)[12]** |  |  |
| 2.60.430 | 1 | 2p84 | A:1-62 | **sheet A: 2=1=5**<br>**sheet B: 3=4** | **1a**<br>**2=1**<br>**3=4**<br>**2(9)[11]** | **1a**<br>**1=5**<br>**3=4**<br>**2(10)[10]** | — | — |
| 2.60.440 | 1 | 2oez | A:1-11,<br>179-245 | **sheet A: 1=4=7=2**<br>**sheet B: 5=6=3** | **2d**<br>**4=7**<br>**5=6**<br>**7(20)[16]** | **2a**<br>**4=7**<br>**6=3**<br>**5+1(16)[14]** | **2d**<br>**7=2**<br>**6=3**<br>**3+1(15)[14]** | — |
| 2.60.450 | 2 | 2r19 | A:28-164 | **sheet A: 1=4=5=8=9=12=13**<br>**sheet B: 2=3=6=7=10=11=14=15** | **2d**<br>**1=4**<br>**2=3**<br>**2(10)[13]** | **2d**<br>**5=8**<br>**6=7**<br>**3+1(13)[13]** | **2d**<br>**9=12**<br>**10=11**<br>**3(11)[12]** | **1b**<br>**12=13**<br>**14=15**<br>**3(11)[12]** |
| 2.60.470 | 1 | 2qts | A: 74-96,<br>A: 165-225 | **sheet A:** 12=1<br>**sheet A:** 11=10=12=7=3=4<br>**sheet B:** 9=8=2=6=5<br>No SubS for strands # 1 and 2 | **1b**<br>11=10<br>9=8<br>4 (19)[20] | **2a**<br>7=3<br>2=6<br>6(16)[13] | **1a**<br>3=4<br>6=5<br>4(12)[12] | **2d**<br>7=3<br>8=2<br>5(16)[13] |
| 2.60.480 | 1 | 3g74 | A:0-90 | **sheet A:** 3=1=5<br>**sheet B:** 4=6=2<br>strands #1 and 5 are parallel<br>strands #2 and 6 are parallel | **2d**<br>3=1<br>4=6<br>6+1(14)[13] | **2a**<br>1=5<br>6=2<br>4(14)[13] | **2a\***<br>3=1<br>6=2<br>6(15)[13] |  |
| 2.60.490 | 1 | 3h9v | A: 59-332 | **sheet A:** 5=7<br>**sheet A:** 1=5=6<br>**sheet B:** 3=4=2=11=8=9<br>**sheet B:** 11=10 | **2d**<br>1=5<br>4=2<br>3(17)[19] | **1b**<br>5=6/7<br>8=9<br>7(18)[18] |  |  |
| 2.60.500 | 1 | 1ywm | A: 63-160 | **sheet A:** 1=4<br>**sheet B:** 2=3=8=7=5=6<br>No SubS for strands # 4 and 5 | **2d**<br>1=4<br>2=3<br>3 (11) [9] | — | — | — |
| 2.60.510 | 2 | 3tcq | A: 204-308 | **sheet A:** 2=1=6<br>**sheet B:** 3=4=5 | **1a**<br>2=1<br>3=4<br>4+1(14)[12] | **1a\***<br>1=6<br>4=5<br>5( 15)[13] | — | — |
| 2.60.520 | 1 | 1zru | A:2-140 | **sheet A:** 1=4=3=2<br>**sheet B:** 6=5=7=8 | **1b**<br>4=3<br>6=5<br>7(20) [19] | — | — | — |

<u>Substructure Type 2</u> has two pairs of consecutive strands: *i* and *i*+1, and *j* and *j*+1 (Fig. 2). Two consecutive strands in each pair are located in different beta-sheets, e.g. strand *i* is located in one sheet and strand *i+1* is in the other sheet. The strands from the different pairs are neighbors in the same sheet and share backbone hydrogen bond contacts, e.g. strand *i* and *j* are located next to each other in sheet A and strands *i+1* and *j+1* - in sheet B. The four possible variants of substructures type 2 – subtypes 2a, 2b, 2c, and 2d - are shown in Fig 2a-d, respectively. Substructure 2a is found in 23 folds, 2b – in 3 folds, 2c – in 8, folds, 2d - in 32 folds (Table 1, column “SubS 1-4”).

**Figure 2.**
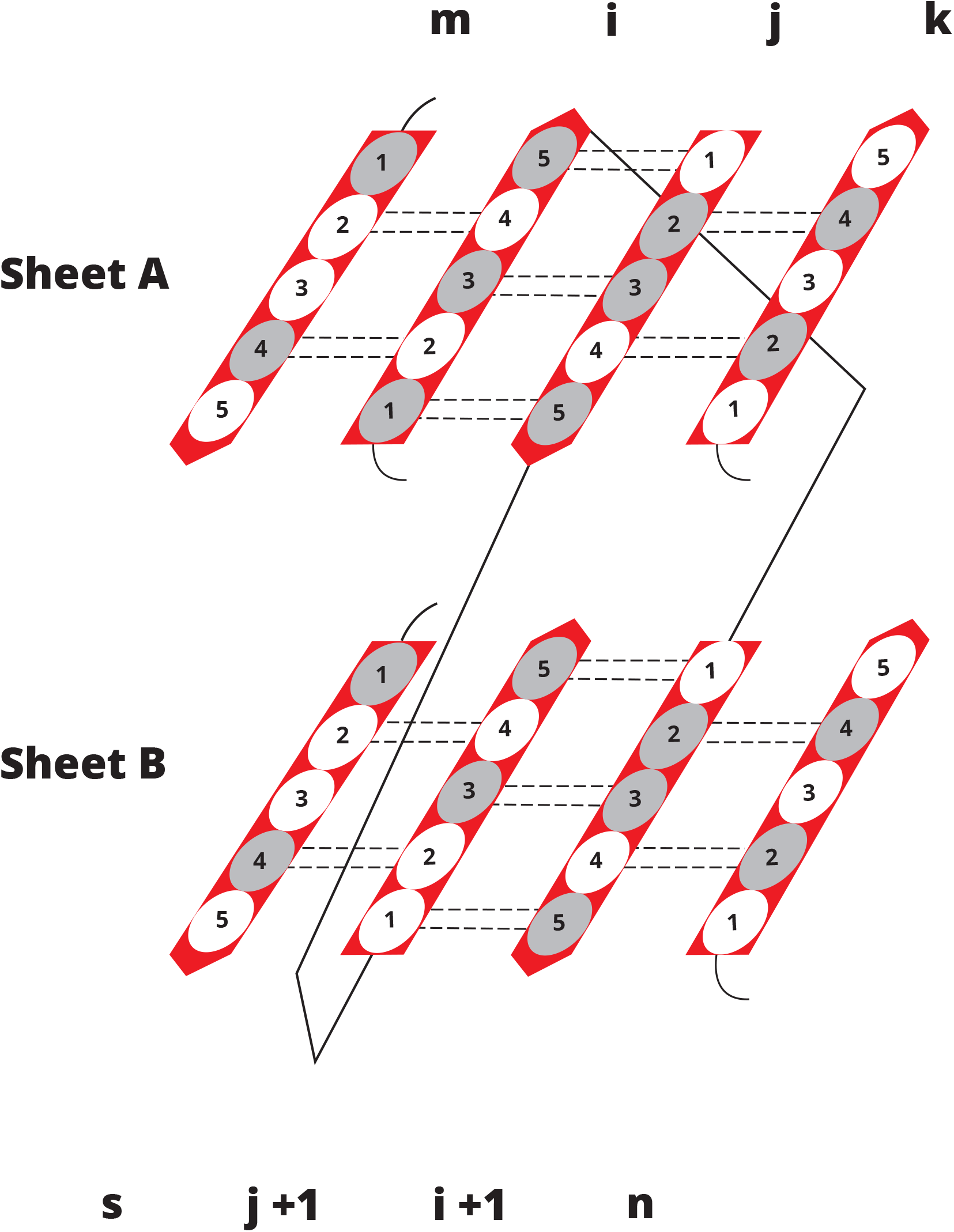
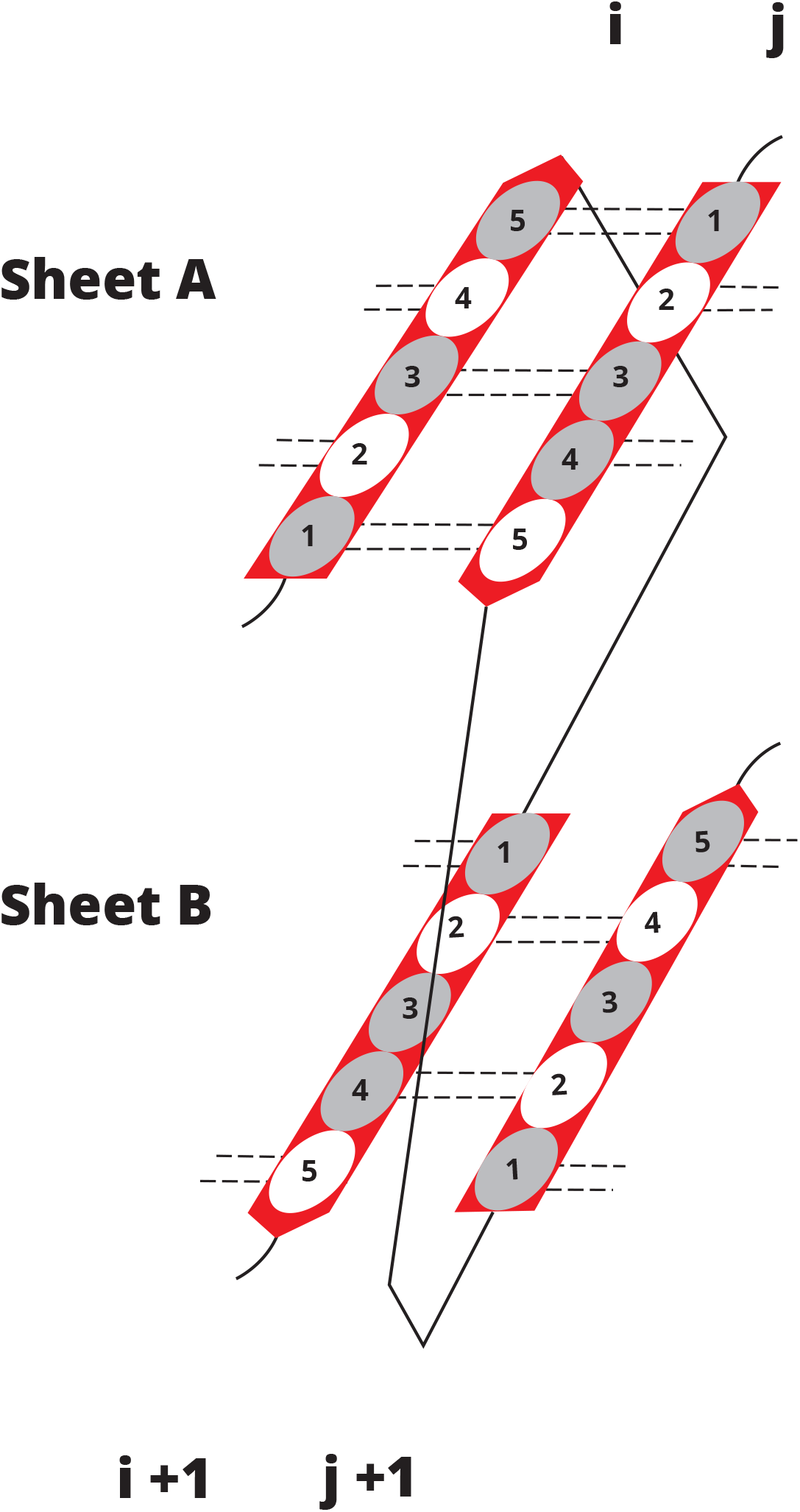
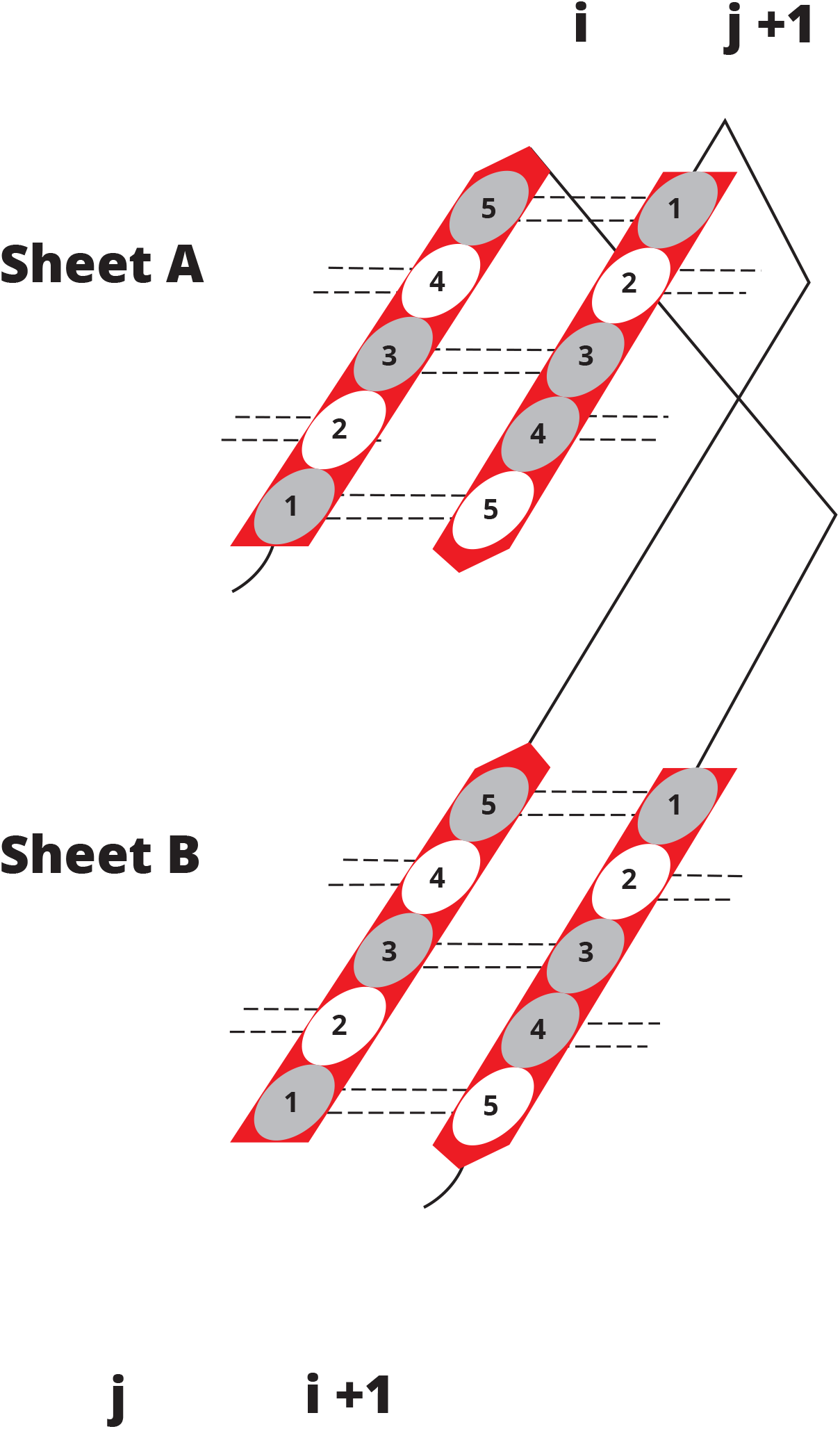
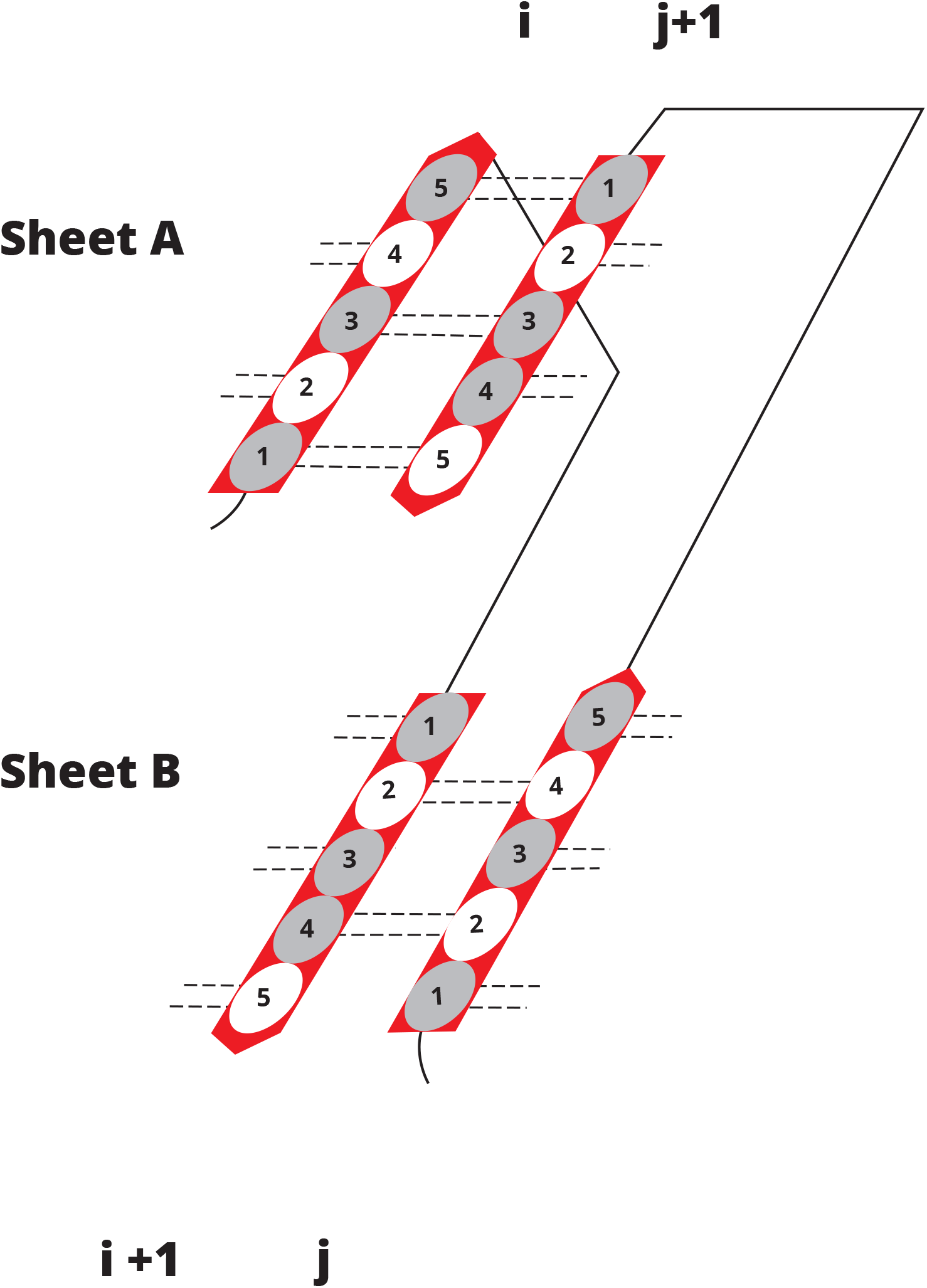
Conserved Substructure type 2. Four variants of Substructures with specific arrangements of strands *i*, *i*+1, *j,* and *j*+1 are shown in 2a, 2b, 2c, and 2d.

#### Substructures in sandwich folds

At least one type 1 or type 2 substructure was found in each domain (Table 1). Most domains contained 2-4 substructures. These restrictions found in the arrangement of substructure strands significantly reduce the number of possible variants of SSS in sandwich domains.

There are two specific characteristics of the substructures. First, the four strands in the substructures form the smallest possible sandwich protein domain. Secondly, SSS substructures describe almost all the observed arrangements of consecutive strands are located in different beta-sheets. There were a total of 166 such pairs of consecutive strands in the examined structures, of which 151 pairs were described by the substructures. For example, in structure 1h8e (Table 1, CATH fold 2.60.15, column ‘SSS domain’), the arrangements of consecutive strands 2 and 3, 5 and 6, 7 and 8, and 9 and 10 are described by substructures 1b, 2c, 1a, 2c, respectively (Table 1, columns ‘SubS 1-4’). One exception in this structure was found for consecutive strands 4 and 5 in two different beta-sheets that are not part of a conserved substructure. Sequential strands that are located in different sheets but are not substructure strands were found mainly at the edges of the beta-sheets.

### Sequence analysis

#### Pairs of contact residues (PCR)

To uncover common sequence characteristics in widely dissimilar proteins from different folds, the concept of ‘Pairs of Contact Residues’ (PCR) was introduced. PCR is defined as the pair of residues that are located opposite each other on two neighboring strands within one beta-sheet. For the pair of antiparallel strands *i*-2 and *i*-1 shown in Fig. 1a, examples of PCRs are pairs of residues 1 and 5, 2 and 4, etc). If there is a ‘bulge’ in a strand, then residues that make up the bulge are not eligible for PCRs. For example, in table 2a (structure 1h8e, fold 2.60.15) residue F28, which make up the bulge in strand #2 (sheet A) has no opposite residue in strand # 1. Thus, the total number of PCRs in pair of strands #1 and 2 is seven, as shown in Table 2a, the column “Characteristics pair of strands”, in square parenthesis.

**Table 2a:**
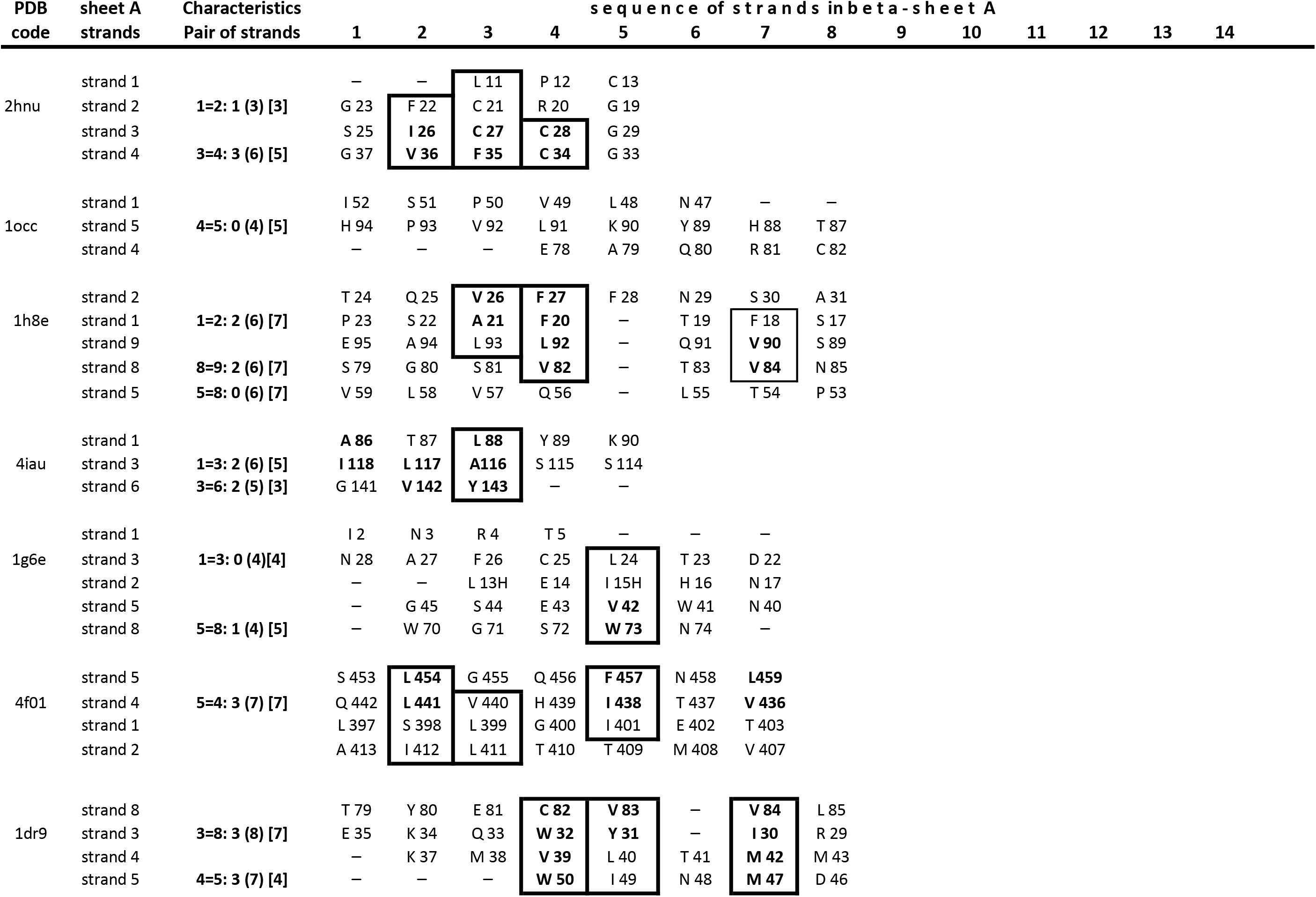
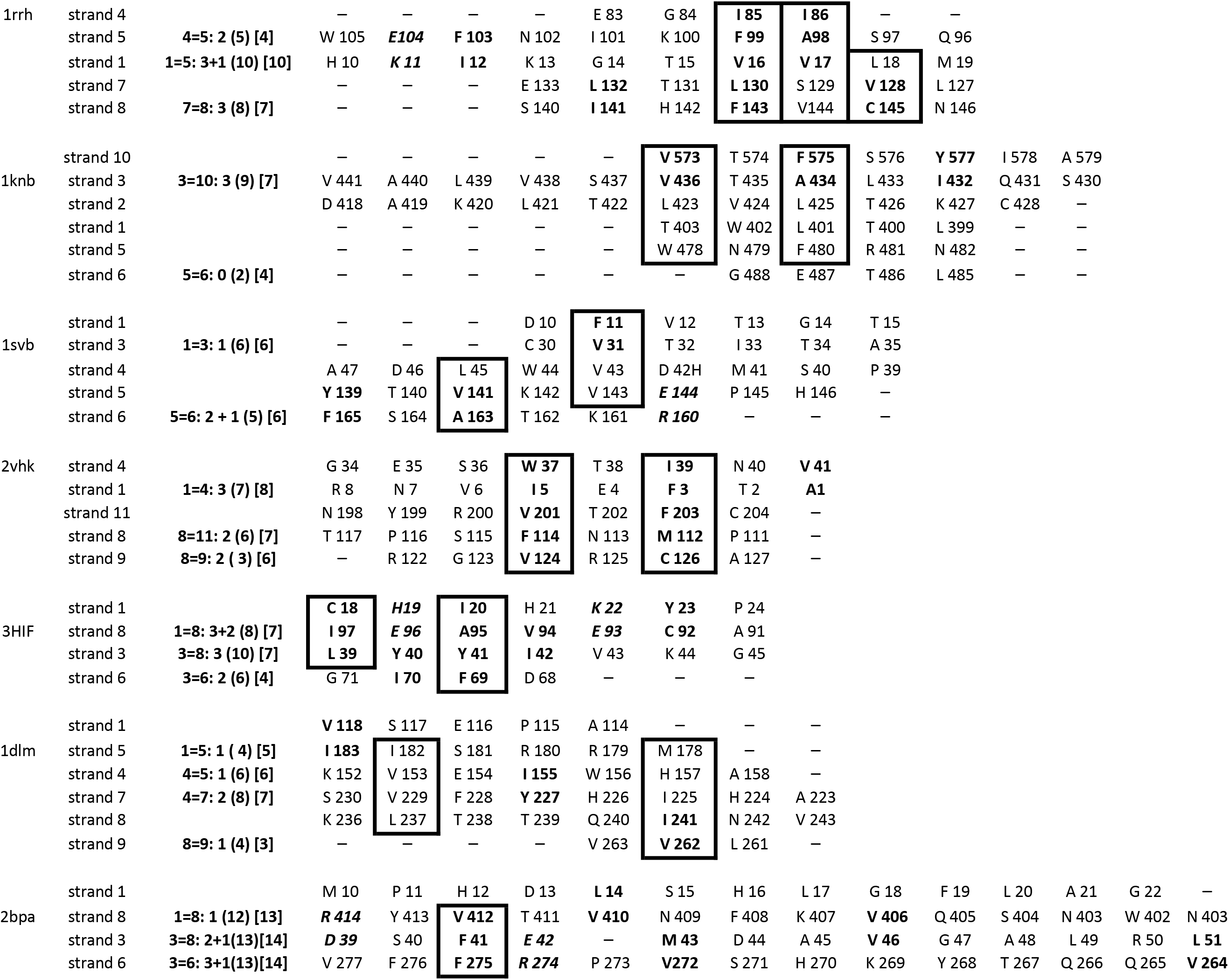
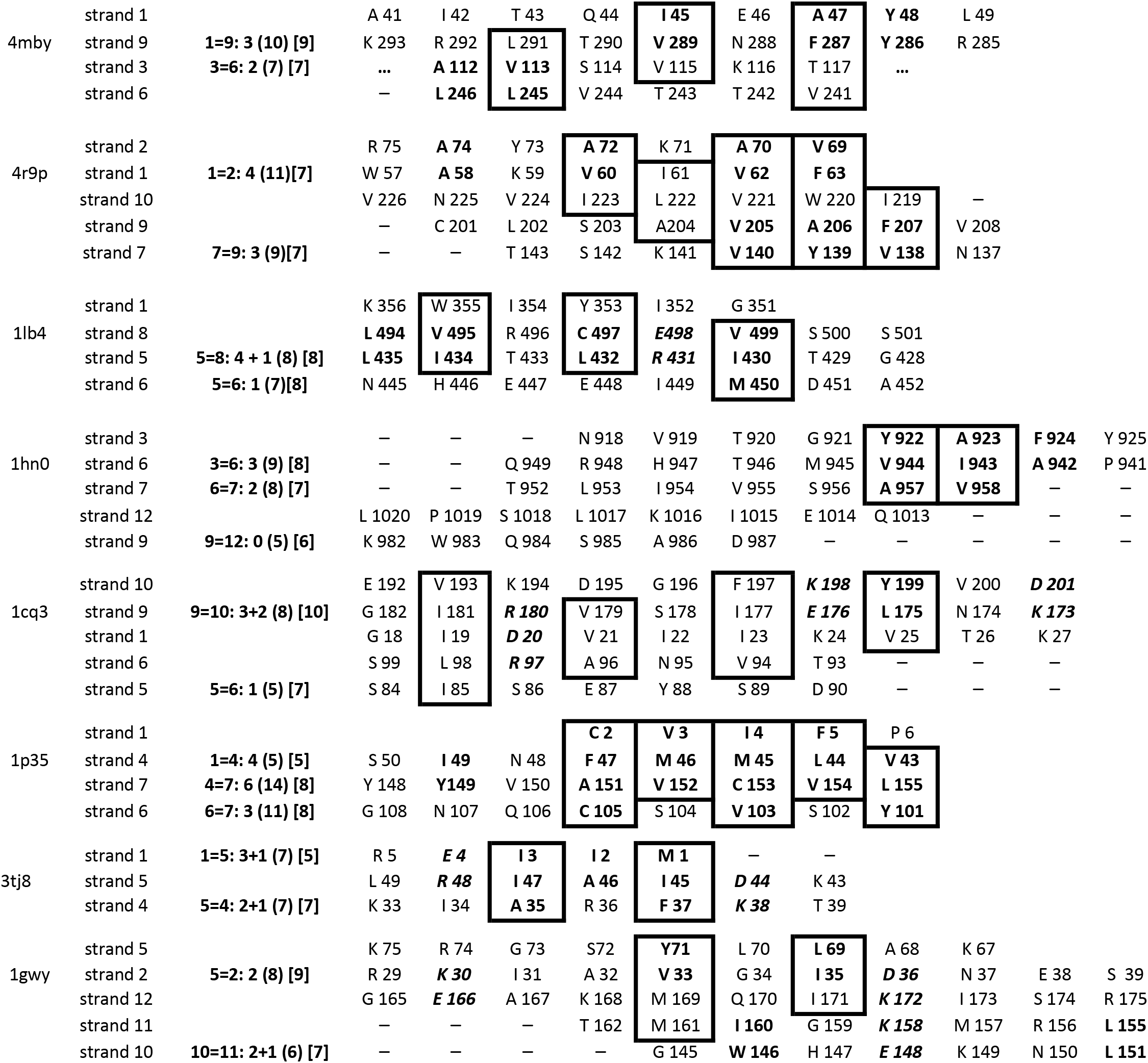
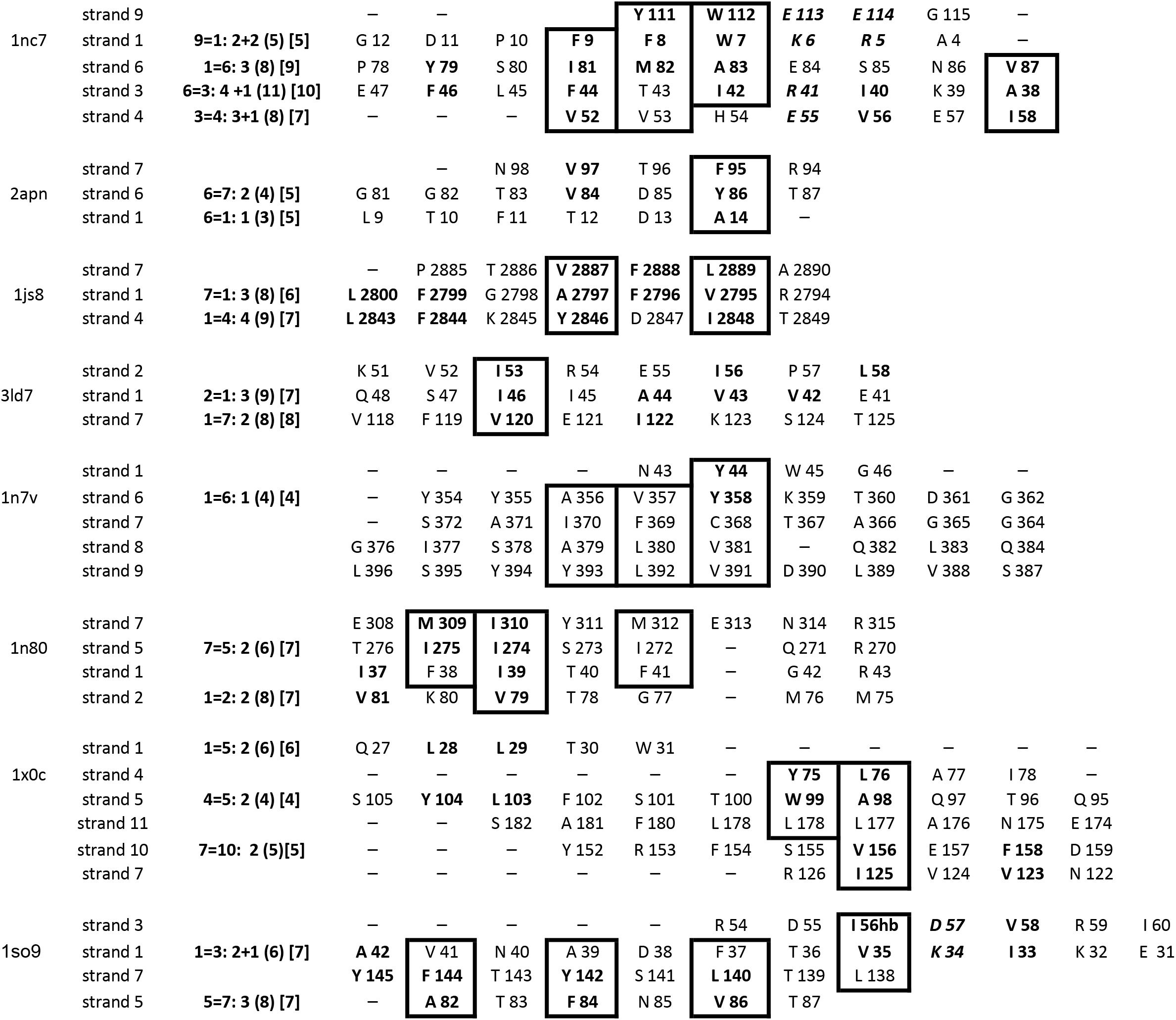
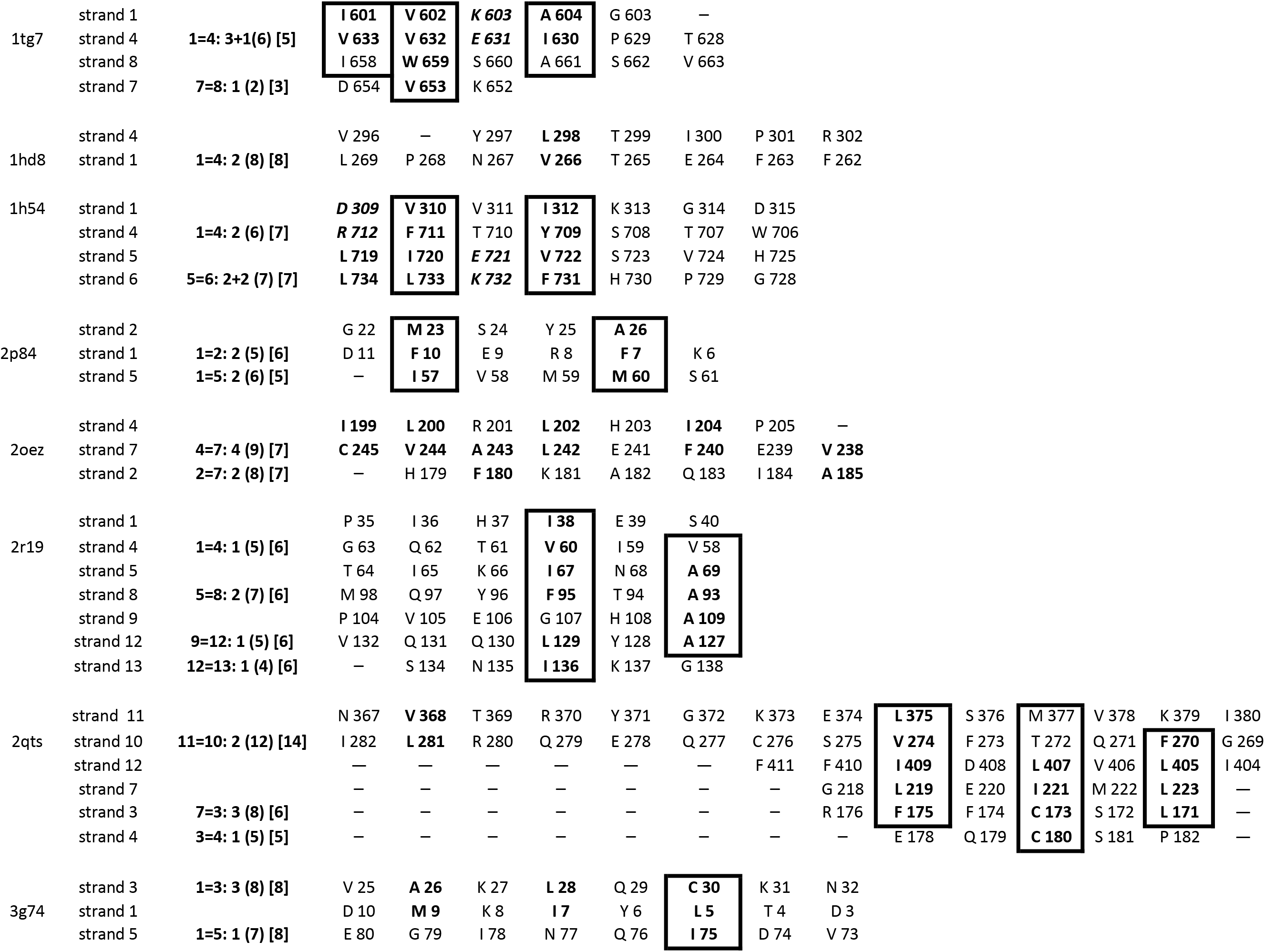
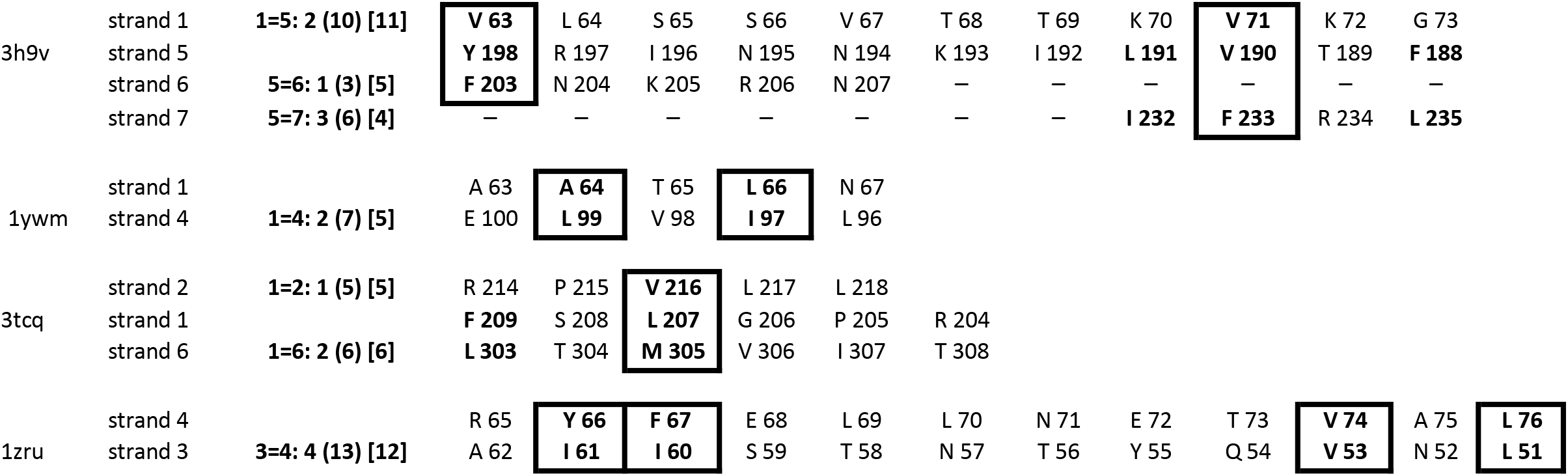
Sandwich domains in 42 different folds. Sequence of strands. Characteristics of pair of strands in beta-sheet A. Columns: **PDB code**: see the legend for this column in Table 1; **Sheet A strands**: strands that form beta-sheet A; **Characteristics Pair of strands**: for pair of strands that form the substructure are presented: 1) the total number of PHR in two pairs of hydrogen-bonded strands (for example, in structure 2hnu the number of PHR is 1 for pair of strands #1 and 2); 2) the number of hydrophobic residues is shown in round brackets (for example, in structure 2hnu the number of hydrophobic residues is 3 for pair of strands #1 and 2); 3) the total number of PCR in square brackets (for example, in structure 2hnu the total number of PCR is 3 for pair of strands #1 and 2); **Sequence of strands in beta-sheets A**: The residues that make up PCRs are shown under each other. The of strands #1 and 2); **Sequence of strands in beta-sheets A**: The residues that make up PCRs are shown under each other. The hydrophobic residues are framed. The pairs of oppositely charged residues are bolded and italicized.

#### Hydrophobic residues in substructures

The total number of hydrophobic residues was equal to, or greater, than the total number of PCRs in about 75% of the substructures. In 18% of substructures, the total number of hydrophobic residues is one less than the total number of PCRs (Table 1, columns “SubS 1-4”). These observations highlight the high proportion of hydrophobic residues in the substructure strands.

A subset of PCRs in which both residues are hydrophobic was called ‘Pairs of Hydrophobic Residues’ (PHRs). The total numbers of PHRs in all substructures of the sandwich domains varied from about 25% to 50% of the total number of PCRs. In 88% of the substructures, the number of PHRs varied from 3 to 7 (Table 1, columns “SubS 1-4”).

Additional evidence for the importance of specific distribution hydrophobic residues to structure formation derives from the location of residues in strands of beta sheets A and B (see columns sequences of strands in Table 2a and 2b). Residues forming PHR are located directly one below each other, which corresponds to the actual location of the strands relative to each other in the beta-sheet. The representations of residues showed that a significant number of hydrophobic residues are not randomly distributed in beta sheets, but most are in columns with predominantly hydrophobic residues in all substructures (see framed PHRs in the columns ‘sequences of strands in beta-sheets A and B” in Table 2a and 2b).

**Table 2b:**
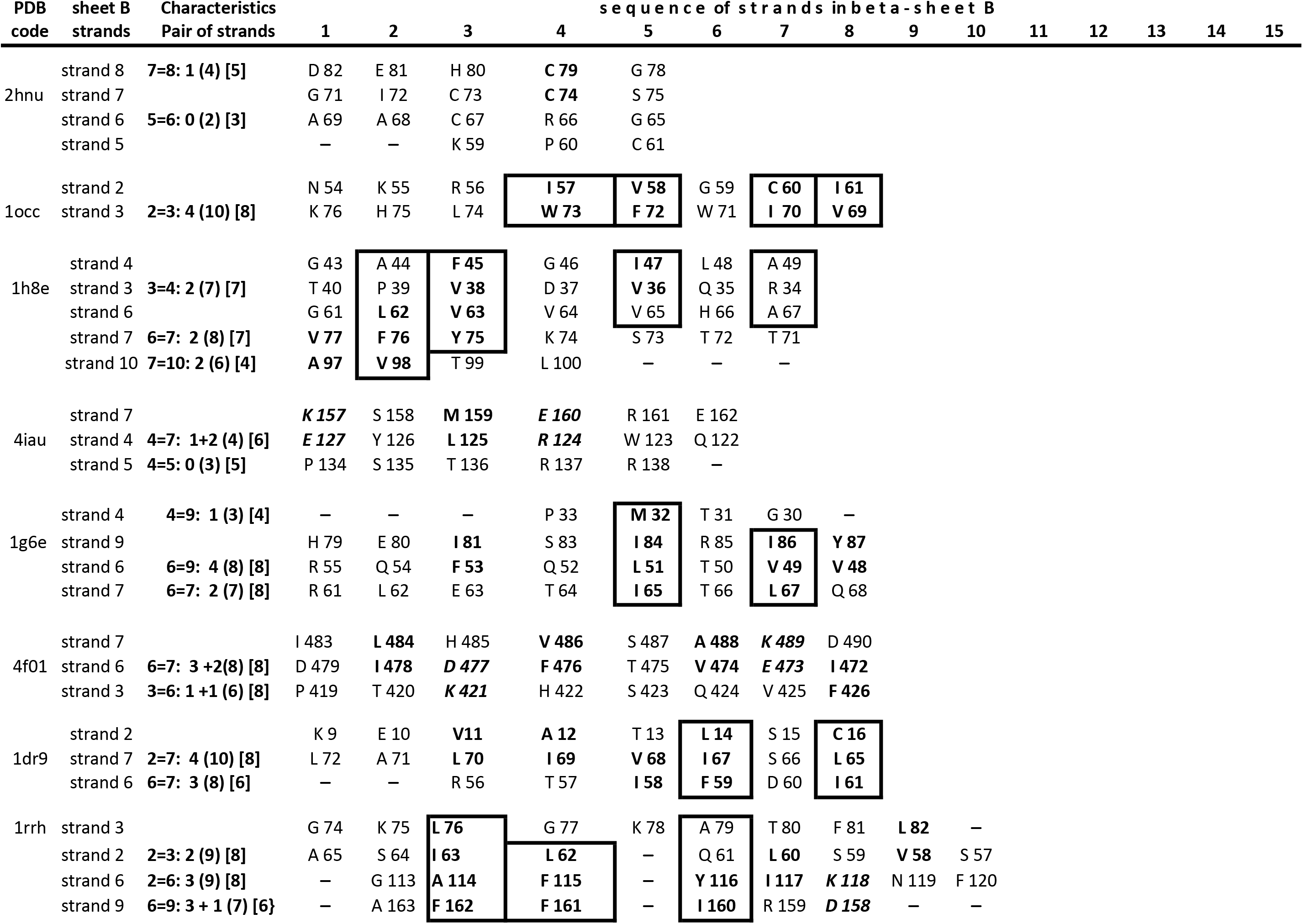
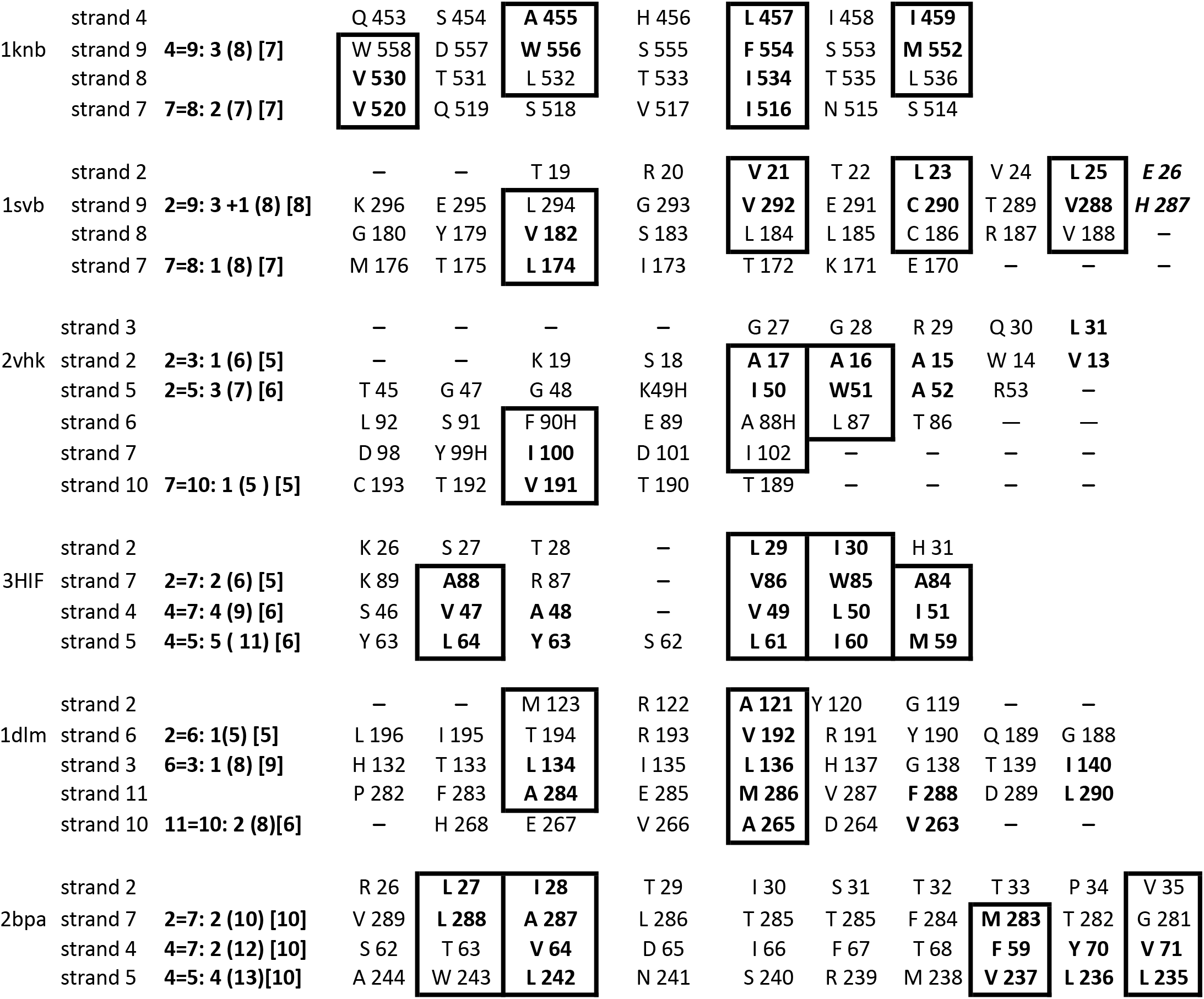
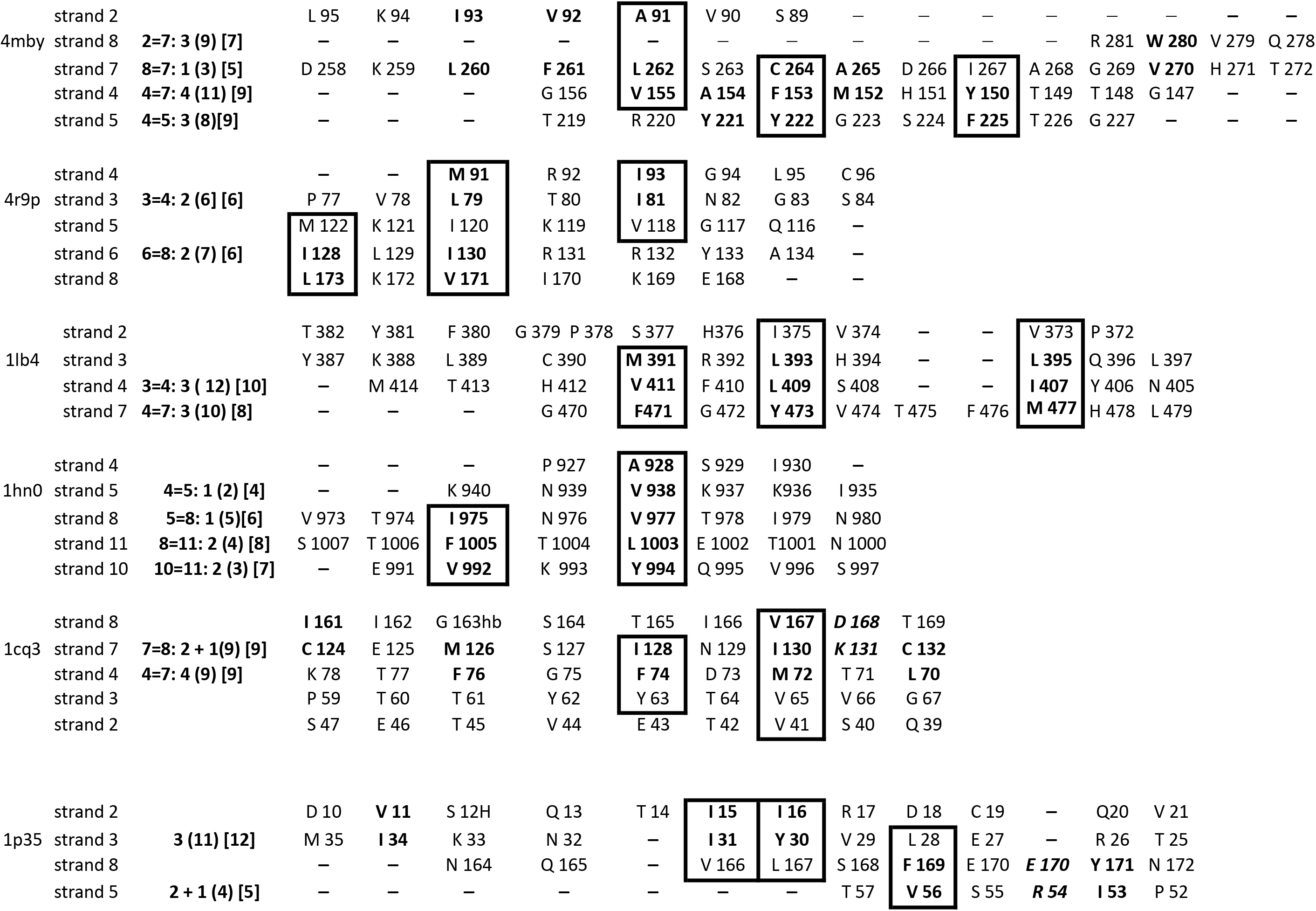
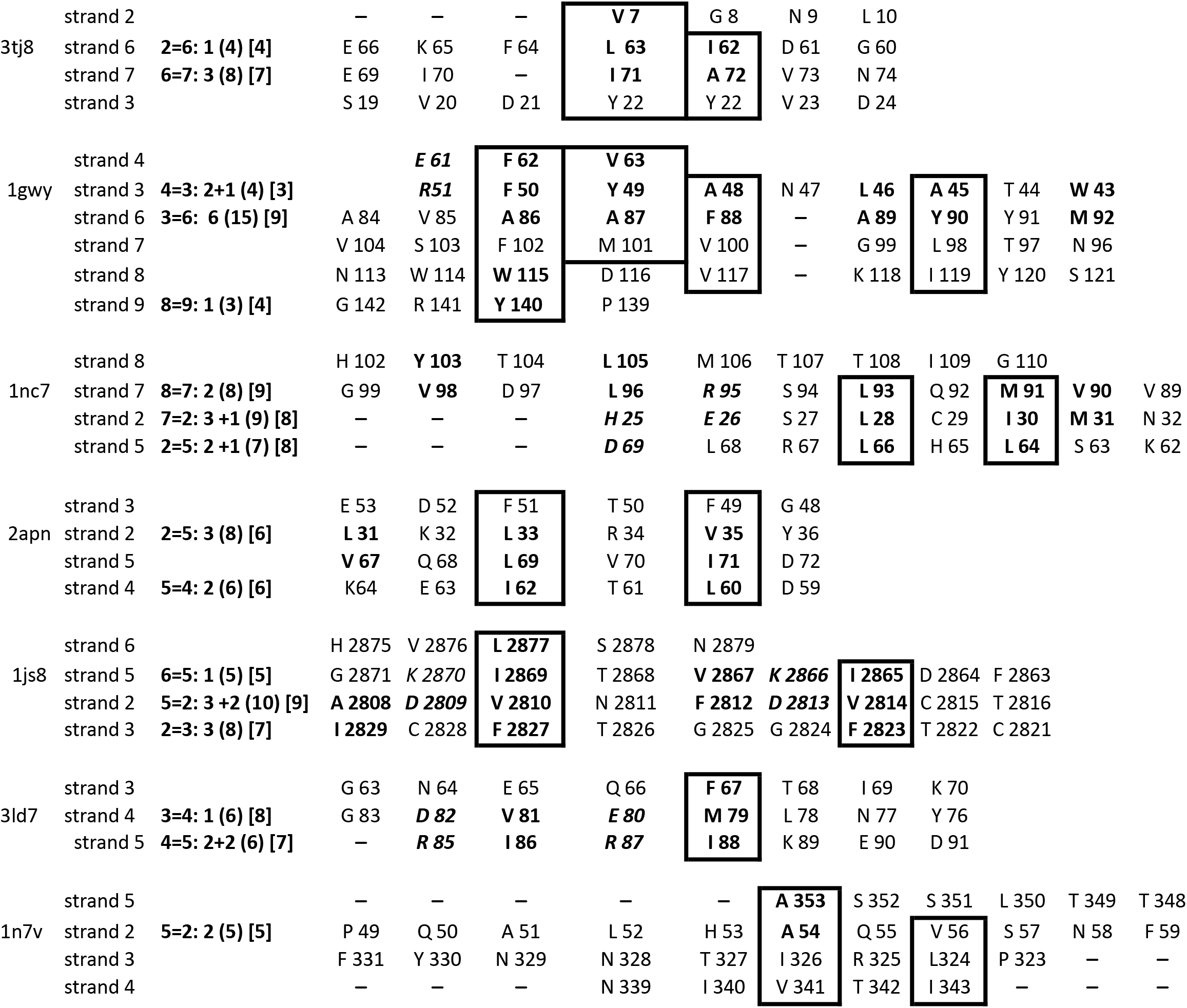
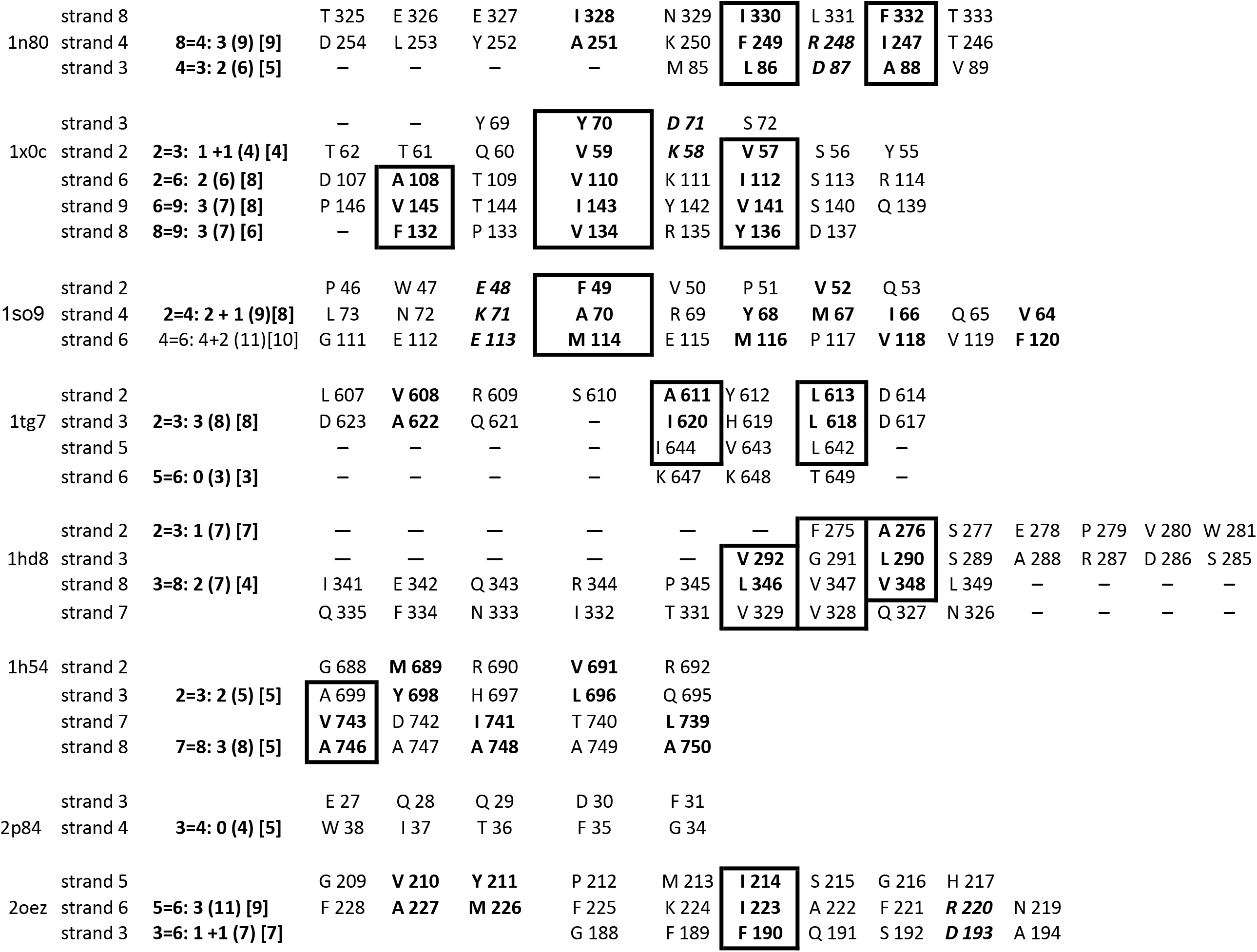
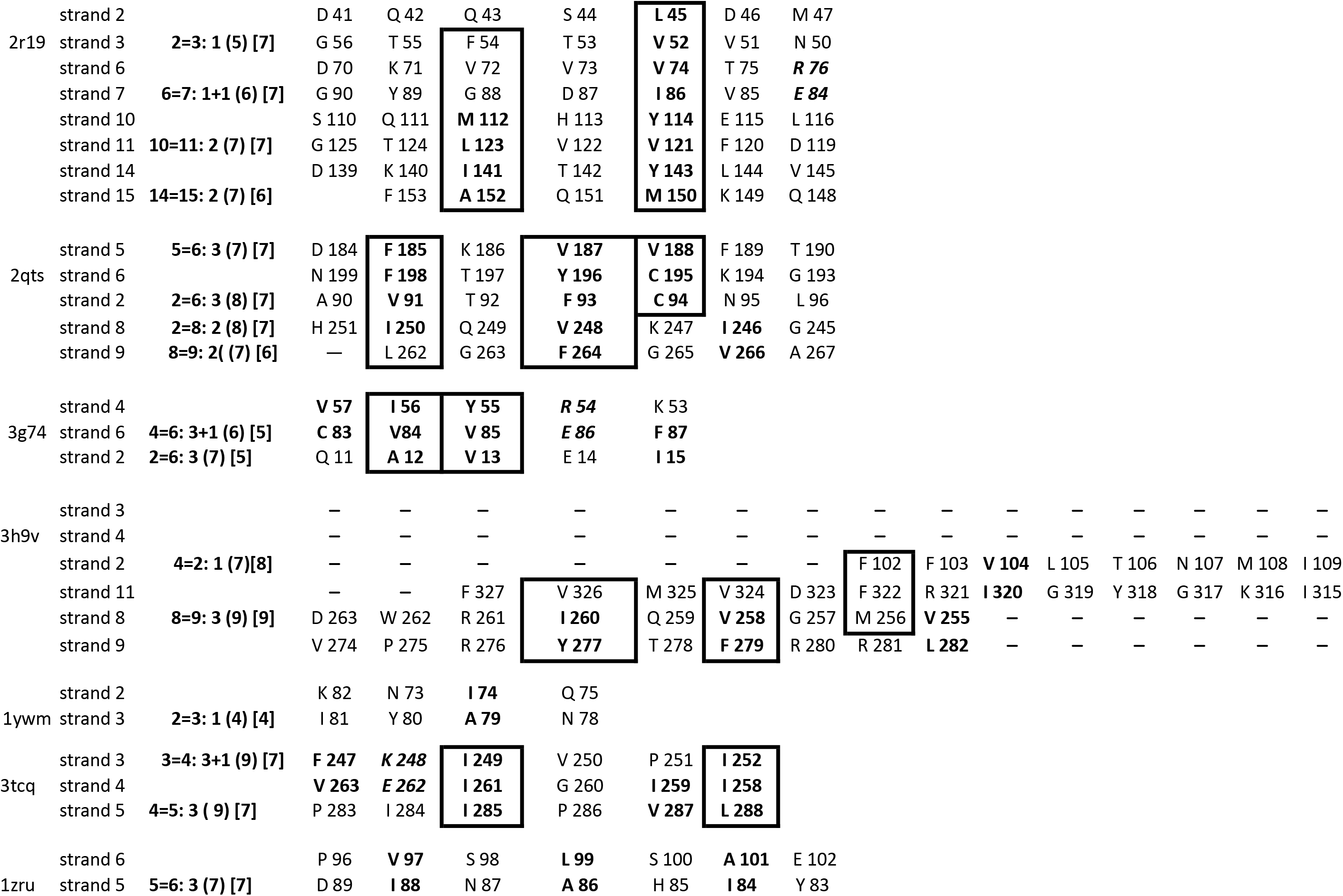
Sandwich domains in 42 different folds. Sequence of strands. Characteristics of pair of strands in beta-sheet B. The legend is similar to the legend of Table 2a

#### Pairs of charged residues

Protein folding initiation sites contain not only hydrophobic amino acids but also charged residues.^18^ Thus, pairs of oppositely charged residues were taken into account in order to uncover the common sequence elements of sandwich-like proteins. The pairs of charged residues were found in almost 20% of neighboring substructure strands in the same sheet. (Table 2a, b in the columns “Sequences of strands in beta-sheets A and B”).

#### Inter-sheet contacts of hydrophobic residues within substructures

A Pair of Hydrophobic Residues that form inter-sheets contacts was called PHR-IC. The numbers of PHR-ICs for each substructure are shown in square brackets in the column “Intersheet Contacts” (Tables 3 and 4). At least one PHR-IC was found in all substructures, and most substructures have 3 and more PHR-ICs. The residues of PHR-IC are shown in columns “Residue-residue intersheet contacts (PHR-IC)” in Table # 3 and 4. Hydrophobic residues that comprise PHR determine mutual orientation between pairs of strands in beta sheets, while residues that make up PHR-IC define the relative orientation of beta-sheets. It was found that a residue of at least one PHR is involved in intersheet contact. Thus, the dependence of the formation of PHR in 2D sheet and PHR-IC in 3D sandwich structure on the distribution of hydrophobic residues in amino acid sequences can serve as one of the decisive criteria for the relationship between primary, secondary, and tertiary structures.

**Table 3a.**
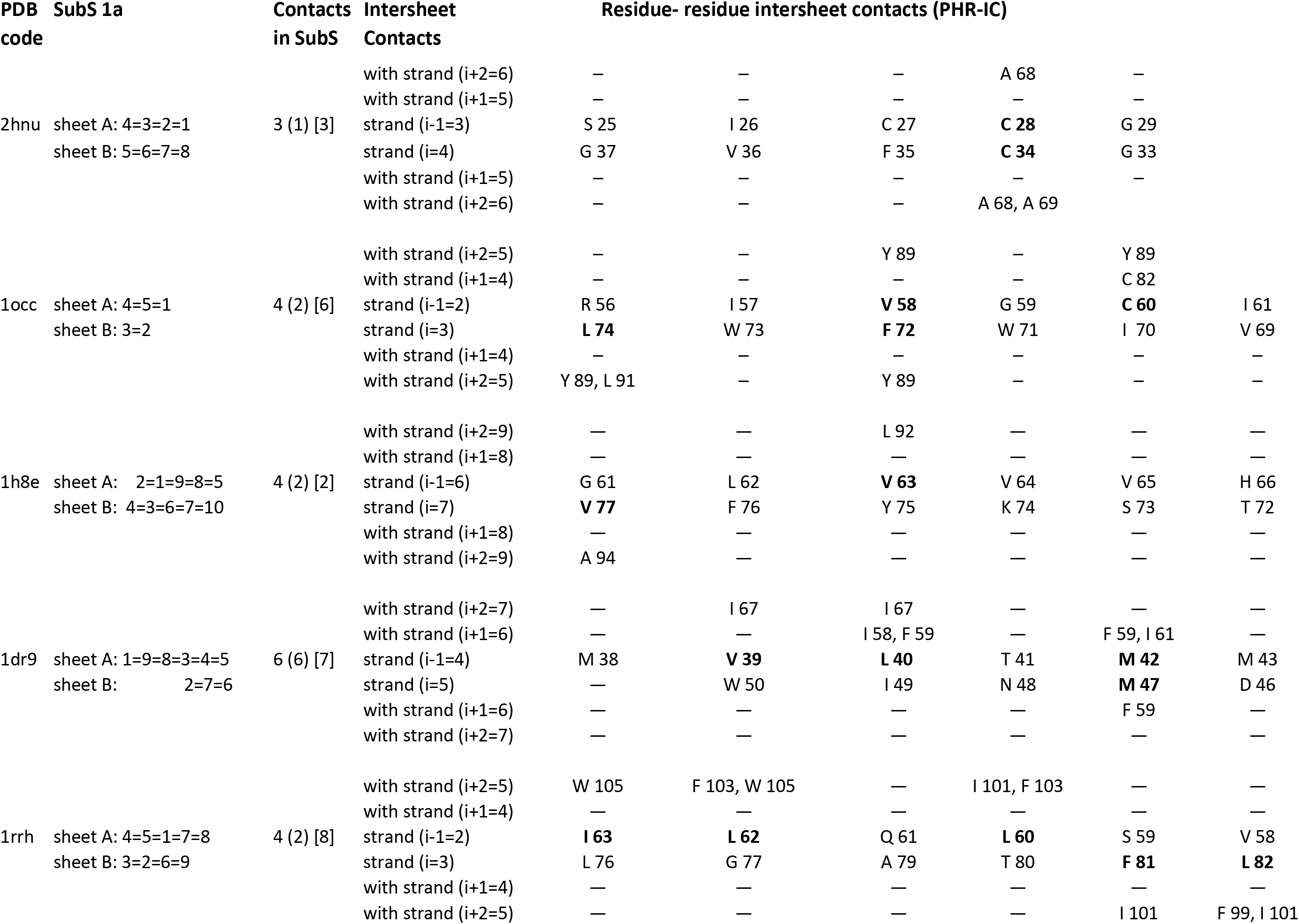
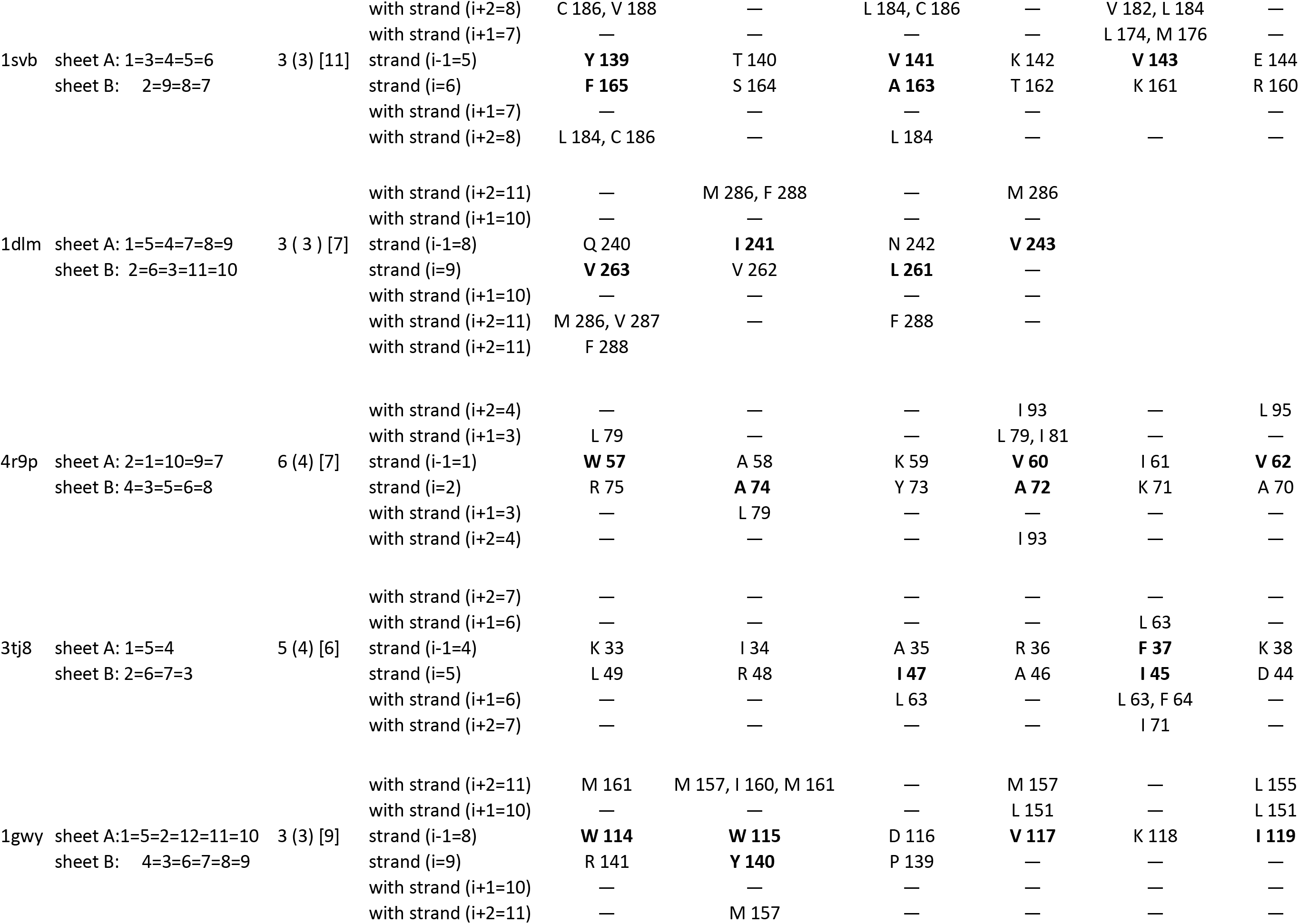
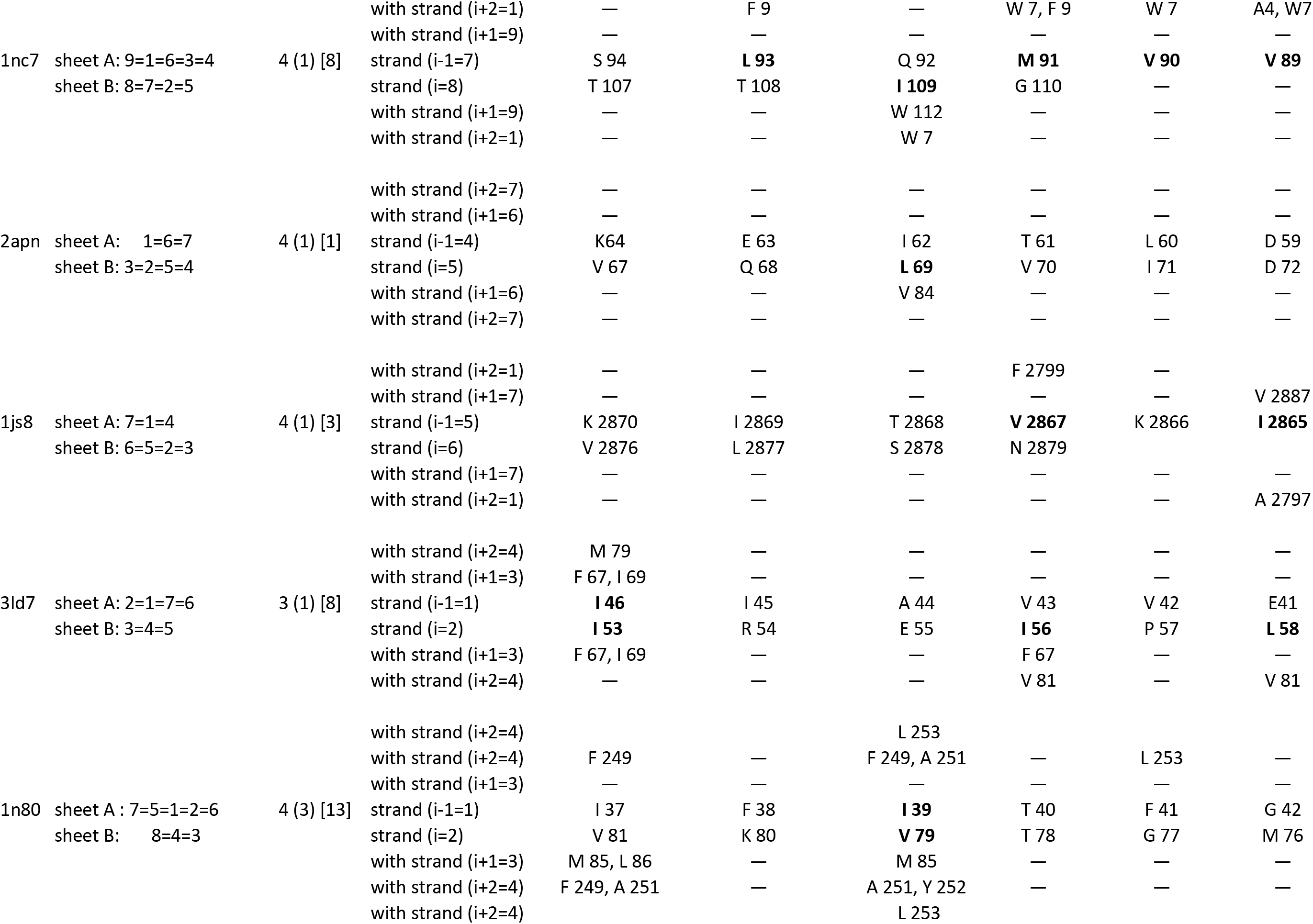
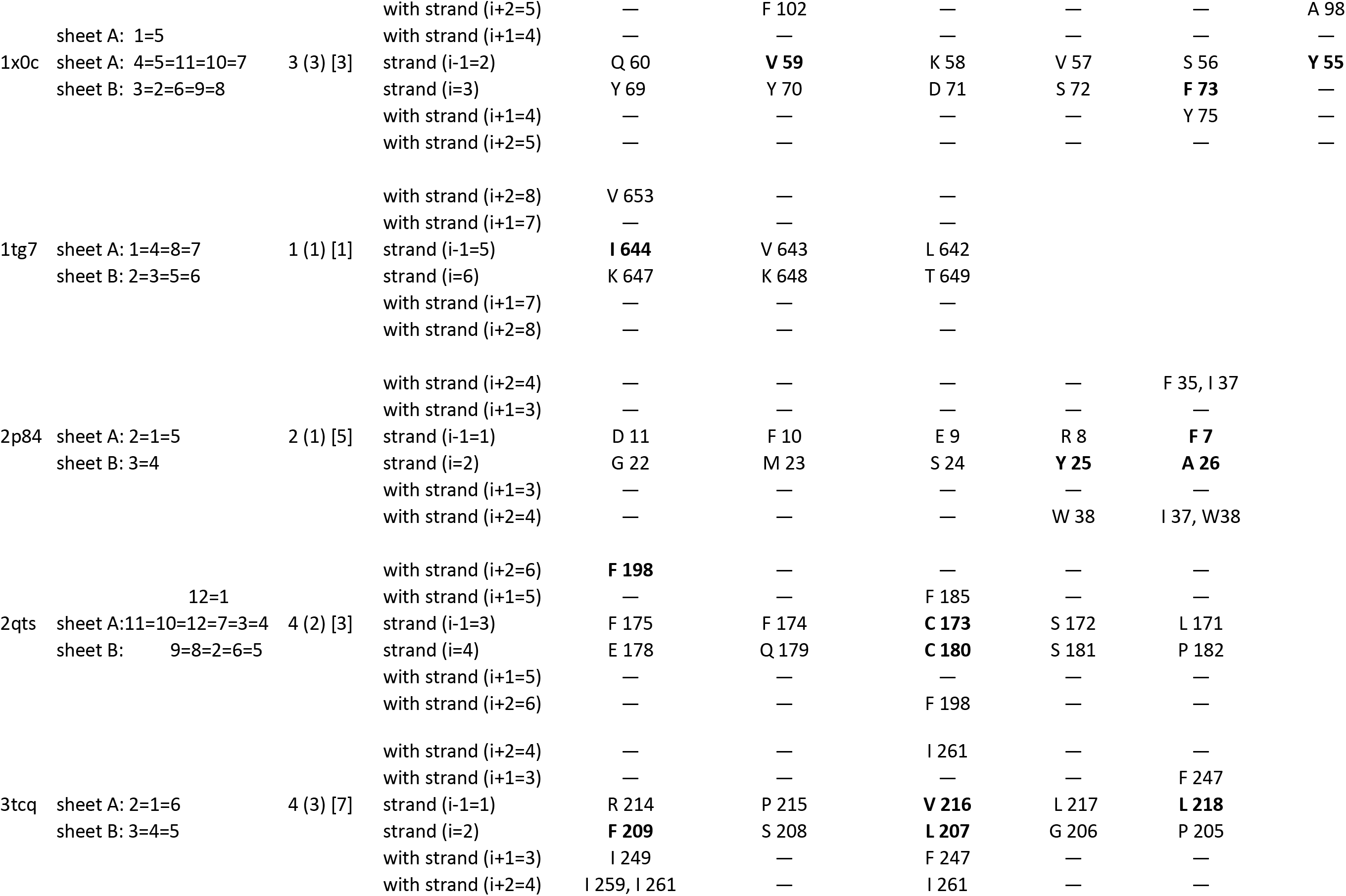
Intersheet contacts of hydrophobic residues in the substructures 1a. Columns: **PDB code** see the legend of Table 1; **SubS 1a**: Supersecondary structure of sandwich domains. SSS of the substructure 1a is indicated by bold and enlarged numbers. **Contacts in SubS** 1) the number of PHRs in the substructure; 2) the number of PHRs whose residues are form intersheet contacts is in round brackets; and 3) the number of PHR-IC in the substructure is in square brackets; **Intersheet Contacts**: between residues in strands i-1 and i in one beta-sheet and residues of strands i+1 and j+1 in another beta sheet; **Residue–residue intersheet contacts (PHR-IC)**: For example, in the structures 1occ residue **L 74** (bolded) in strand i=3 forms two intersheet contacts with residues Y 89 and L 91 in strand (i+2=5). Residue **C 60** in strand (i-1=2) (bolded) forms two PHR-IC with residues C 82 (i+1=4) and Y 89 in strand (i+2=5).

**Table 3b.**
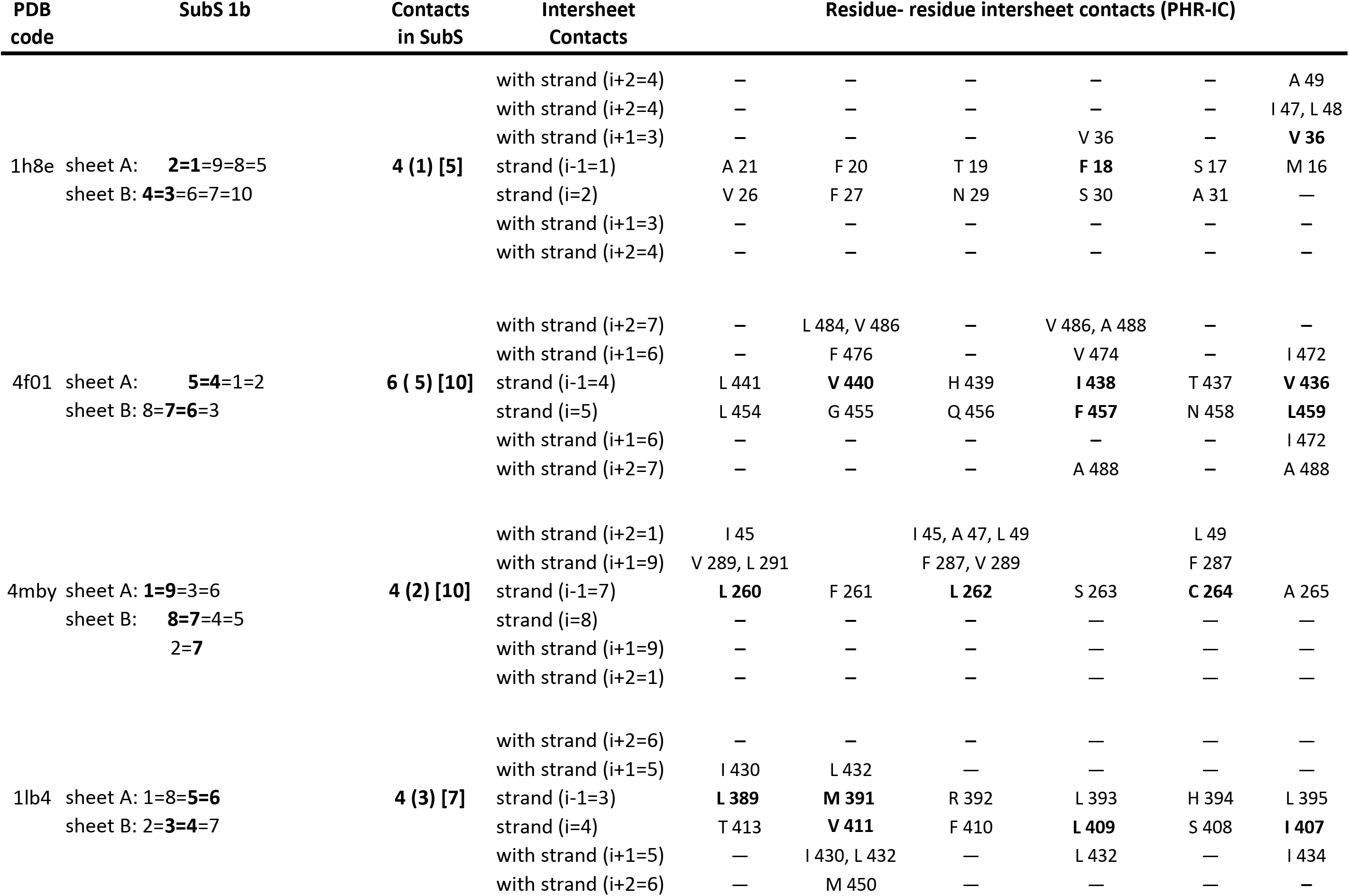
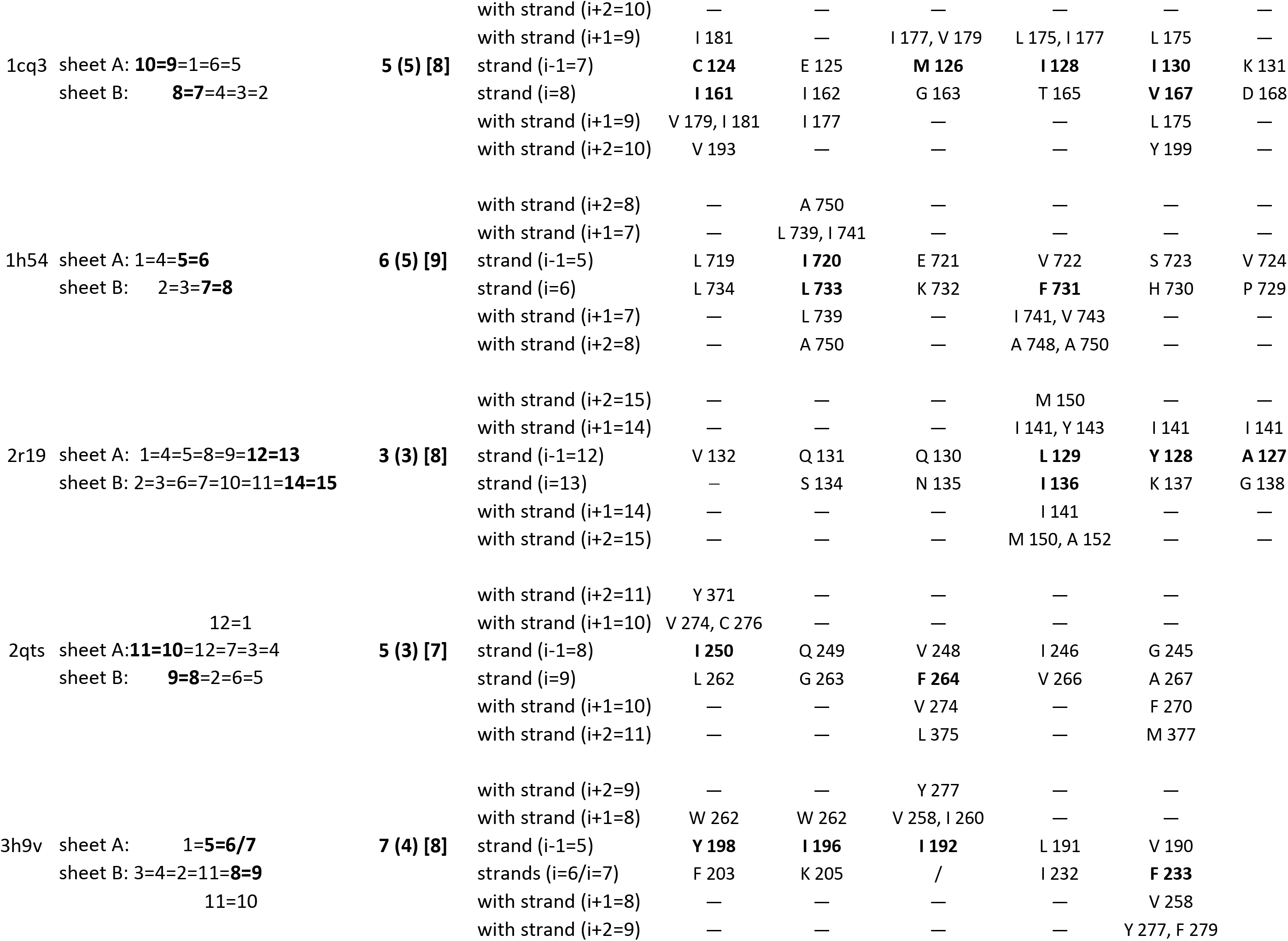
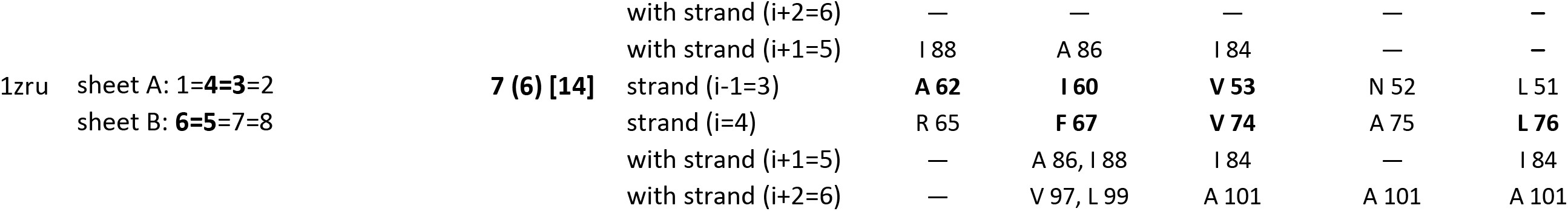
Intersheet contacts of hydrophobic residues in the substructures 1b. The legend is similar to the legend of Table 3a

| PDB<br>code | SubS 1b | Contacts<br>in SubS | Intersheet<br>Contacts | Residue- residue intersheet contacts (PHR-IC) |  |  |  |  |  |
| --- | --- | --- | --- | --- | --- | --- | --- | --- | --- |
| 1h8e | sheet A: 2=1=9=8=5<br>sheet B: 4=3=6=7=10 | 4 (1) [5] | with strand (i+2=4) | — | — | — | — | — | A 49 |
|  |  |  | with strand (i+2=4) | — | — | — | — | — | I 47, L 48 |
|  |  |  | with strand (i+1=3) | — | — | — | V 36 | — | V 36 |
|  |  |  | strand (i-1=1) | A 21 | F 20 | T 19 | F 18 | S 17 | M 16 |
|  |  |  | strand (i=2) | V 26 | F 27 | N 29 | S 30 | A 31 | — |
|  |  |  | with strand (i+1=3) | — | — | — | — | — | — |
|  |  |  | with strand (i+2=4) | — | — | — | — | — | — |
| 4f01 | sheet A: 5=4=1=2<br>sheet B: 8=7=6=3 | 6 ( 5) [10] | with strand (i+2=7) | — | L 484, V 486 | — | V 486, A 488 | — | — |
|  |  |  | with strand (i+1=6) | — | F 476 | — | V 474 | — | I 472 |
|  |  |  | strand (i-1=4) | L 441 | V 440 | H 439 | I 438 | T 437 | V 436 |
|  |  |  | strand (i=5) | L 454 | G 455 | Q 456 | F 457 | N 458 | L459 |
|  |  |  | with strand (i+1=6) | — | — | — | — | — | I 472 |
|  |  |  | with strand (i+2=7) | — | — | — | A 488 | — | A 488 |
| 4mby | sheet A: 1=9=3=6<br>sheet B: 8=7=4=5<br>2=7 | 4 (2) [10] | with strand (i+2=1) | I 45 |  | I 45, A 47, L 49 |  | L 49 |  |
|  |  |  | with strand (i+1=9) | V 289, L 291 |  | F 287, V 289 |  | F 287 |  |
|  |  |  | strand (i-1=7) | L 260 | F 261 | L 262 | S 263 | C 264 | A 265 |
|  |  |  | strand (i=8) | — | — | — | — | — | — |
|  |  |  | with strand (i+1=9) | — | — | — | — | — | — |
|  |  |  | with strand (i+2=1) | — | — | — | — | — | — |
| 1lb4 | sheet A: 1=8=5=6<br>sheet B: 2=3=4=7 | 4 (3) [7] | with strand (i+2=6) | — | — | — | — | — | — |
|  |  |  | with strand (i+1=5) | I 430 | L 432 | — | — | — | — |
|  |  |  | strand (i-1=3) | L 389 | M 391 | R 392 | L 393 | H 394 | L 395 |
|  |  |  | strand (i=4) | T 413 | V 411 | F 410 | L 409 | S 408 | I 407 |
|  |  |  | with strand (i+1=5) | — | I 430, L 432 | — | L 432 | — | I 434 |
|  |  |  | with strand (i+2=6) | — | M 450 | — | — | — | — |
| 1cq3 | sheet A: **10=9=1=6=5**<br>sheet B: **8=7=4=3=2** | **5 (5) [8]** | with strand (i+2=10) | — | — | — | — | — | — |
|  |  |  | with strand (i+1=9) | I 181 | — | I 177, V 179 | L 175, I 177 | L 175 | — |
|  |  |  | strand (i-1=7) | **C 124** | E 125 | **M 126** | **I 128** | **I 130** | K 131 |
|  |  |  | strand (i=8) | **I 161** | I 162 | G 163 | T 165 | **V 167** | D 168 |
|  |  |  | with strand (i+1=9) | V 179, I 181 | I 177 | — | — | L 175 | — |
|  |  |  | with strand (i+2=10) | V 193 | — | — | — | Y 199 | — |
| 1h54 | sheet A: **1=4=5=6**<br>sheet B: **2=3=7=8** | **6 (5) [9]** | with strand (i+2=8) | — | A 750 | — | — | — | — |
|  |  |  | with strand (i+1=7) | — | L 739, I 741 | — | — | — | — |
|  |  |  | strand (i-1=5) | L 719 | **I 720** | E 721 | V 722 | S 723 | V 724 |
|  |  |  | strand (i=6) | L 734 | **L 733** | K 732 | **F 731** | H 730 | P 729 |
|  |  |  | with strand (i+1=7) | — | L 739 | — | I 741, V 743 | — | — |
|  |  |  | with strand (i+2=8) | — | A 750 | — | A 748, A 750 | — | — |
| 2r19 | sheet A: **1=4=5=8=9=12=13**<br>sheet B: **2=3=6=7=10=11=14=15** | **3 (3) [8]** | with strand (i+2=15) | — | — | — | M 150 | — | — |
|  |  |  | with strand (i+1=14) | — | — | — | I 141, Y 143 | I 141 | I 141 |
|  |  |  | strand (i-1=12) | V 132 | Q 131 | Q 130 | **L 129** | **Y 128** | **A 127** |
|  |  |  | strand (i=13) | — | S 134 | N 135 | **I 136** | K 137 | G 138 |
|  |  |  | with strand (i+1=14) | — | — | — | I 141 | — | — |
|  |  |  | with strand (i+2=15) | — | — | — | M 150, A 152 | — | — |
| 2qts | 12=1<br>sheet A: **11=10=12=7=3=4**<br>sheet B: **9=8=2=6=5** | **5 (3) [7]** | with strand (i+2=11) | Y 371 | — | — | — | — | — |
|  |  |  | with strand (i+1=10) | V 274, C 276 | — | — | — | — | — |
|  |  |  | strand (i-1=8) | **I 250** | Q 249 | V 248 | I 246 | G 245 | — |
|  |  |  | strand (i=9) | L 262 | G 263 | **F 264** | V 266 | A 267 | — |
|  |  |  | with strand (i+1=10) | — | — | V 274 | — | F 270 | — |
|  |  |  | with strand (i+2=11) | — | — | L 375 | — | M 377 | — |
| 3h9v | sheet A: **1=5=6/7**<br>sheet B: **3=4=2=11=8=9**<br>11=10 | **7 (4) [8]** | with strand (i+2=9) | — | — | Y 277 | — | — | — |
|  |  |  | with strand (i+1=8) | W 262 | W 262 | V 258, I 260 | — | — | — |
|  |  |  | strand (i-1=5) | **Y 198** | **I 196** | **I 192** | L 191 | V 190 | — |
|  |  |  | strands (i=6/i=7) | F 203 | K 205 | / | I 232 | **F 233** | — |
|  |  |  | with strand (i+1=8) | — | — | — | — | V 258 | — |
|  |  |  | with strand (i+2=9) | — | — | — | — | Y 277, F 279 | — |

**Table 3b. Intersheet contacts of hydrophobic residues in the substructures 1b.**
|  |  |  |  |  |  |  |  |  |
| --- | --- | --- | --- | --- | --- | --- | --- | --- |
| 1zru | sheet A: 1=**4**=**3**=2<br>sheet B: **6**=**5**=7=8 | **7 (6) [14]** | with strand (i+2=6) | — | — | — | — | — |
|  |  |  | with strand (i+1=5) | I 88 | A 86 | I 84 | — | — |
|  |  |  | strand (i-1=3) | **A 62** | **I 60** | **V 53** | N 52 | L 51 |
|  |  |  | strand (i=4) | R 65 | **F 67** | **V 74** | A 75 | **L 76** |
|  |  |  | with strand (i+1=5) | — | A 86, I 88 | I 84 | — | I 84 |
|  |  |  | with strand (i+2=6) | — | V 97, L 99 | A 101 | A 101 | A 101 |

**Table 4a.**
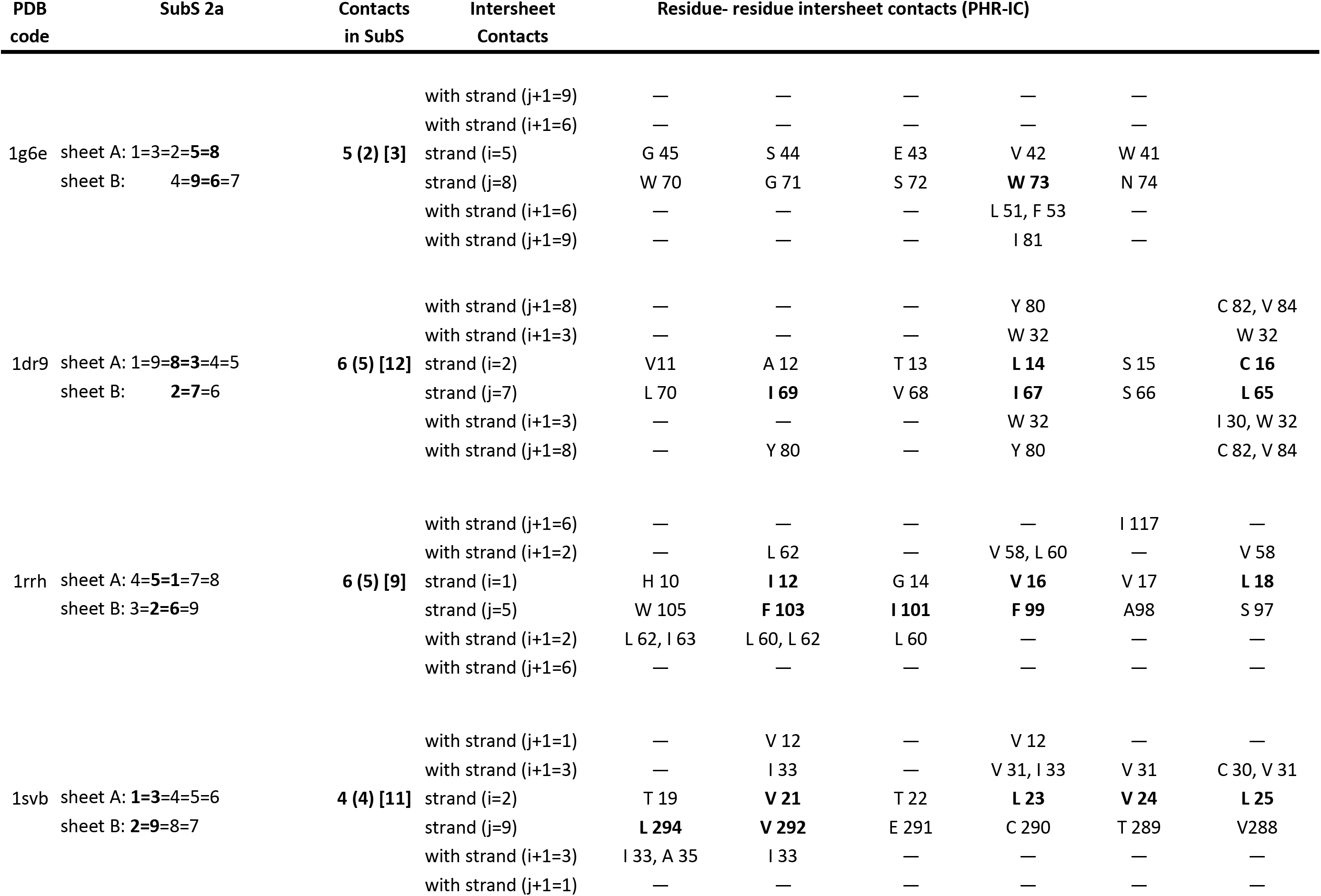
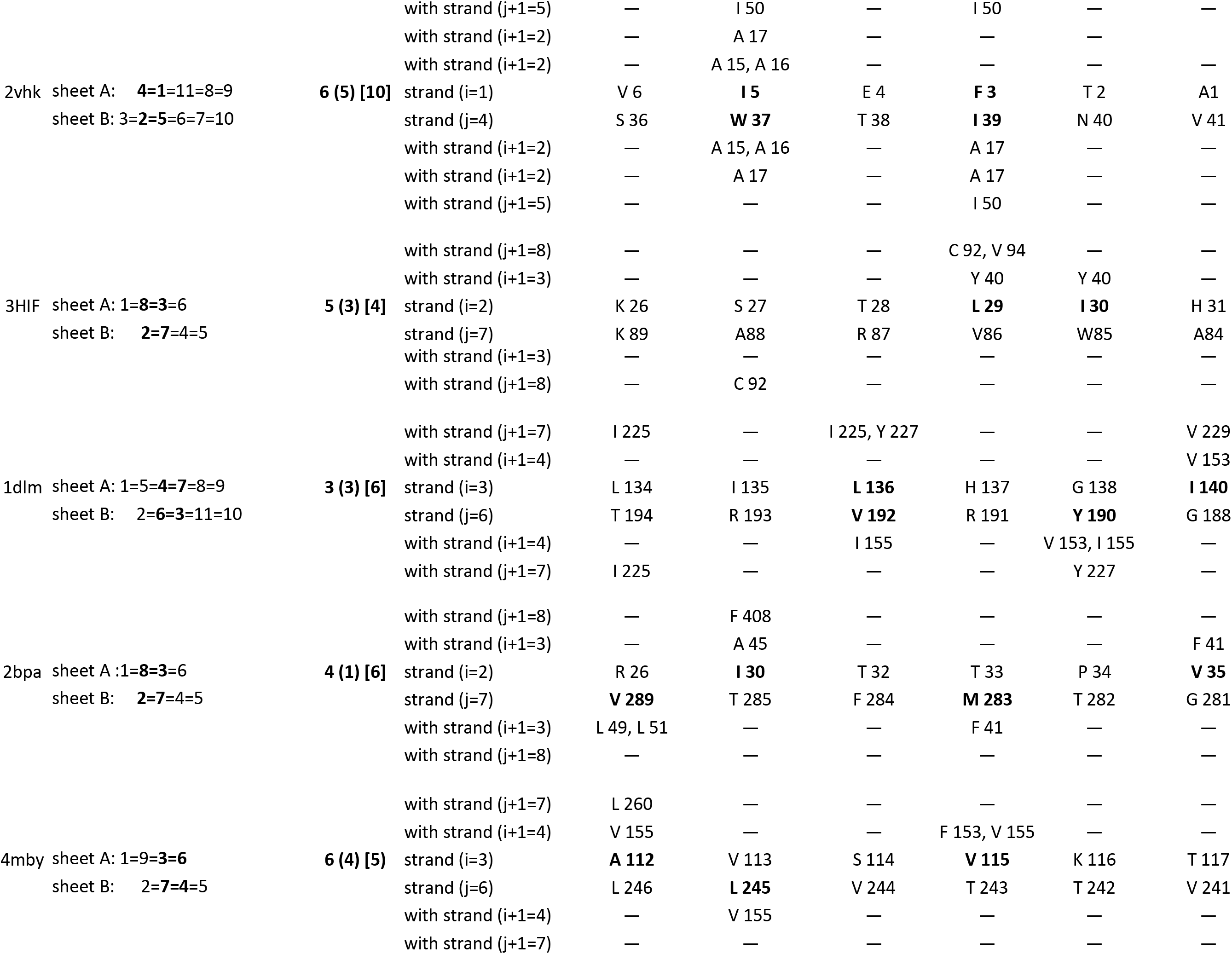
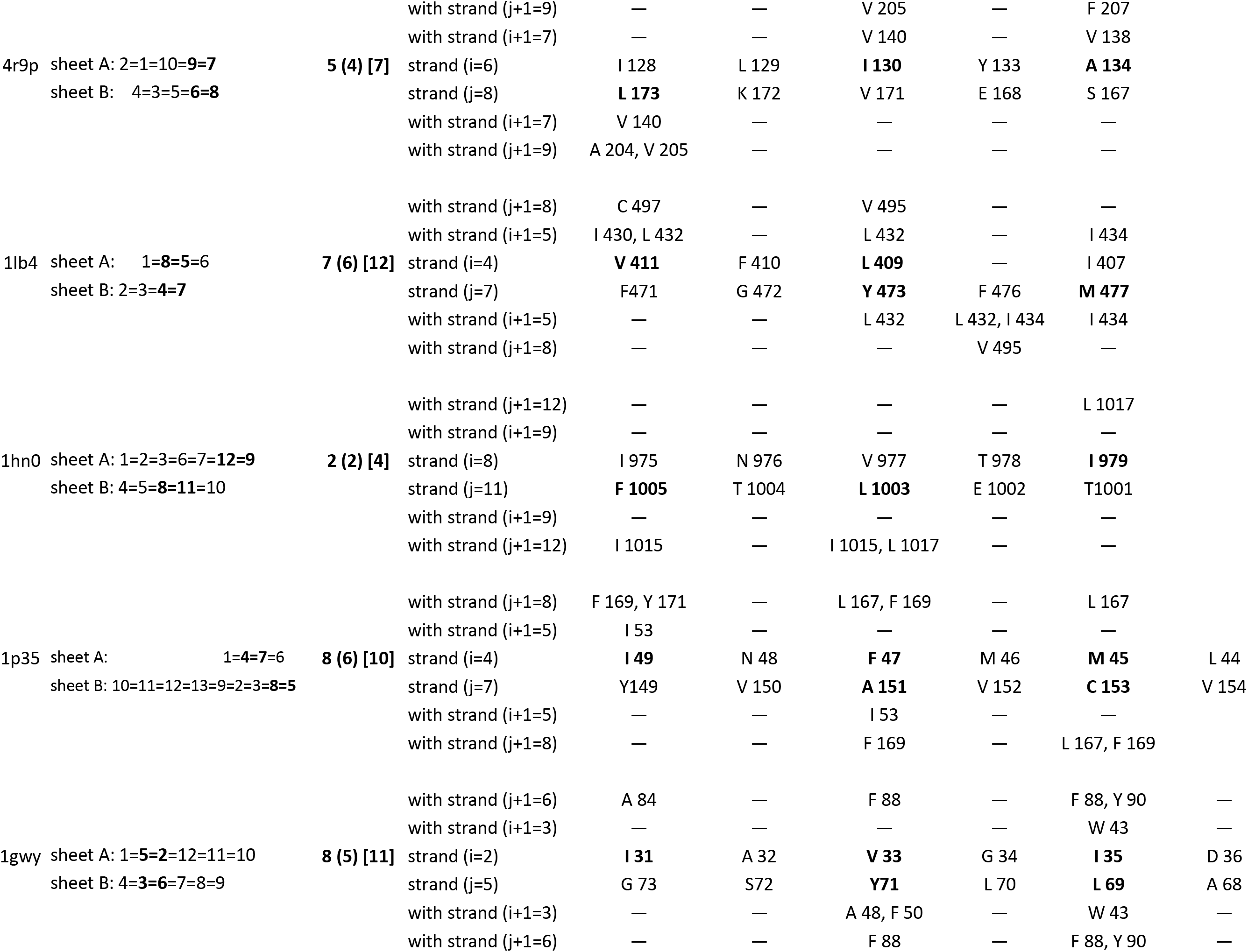
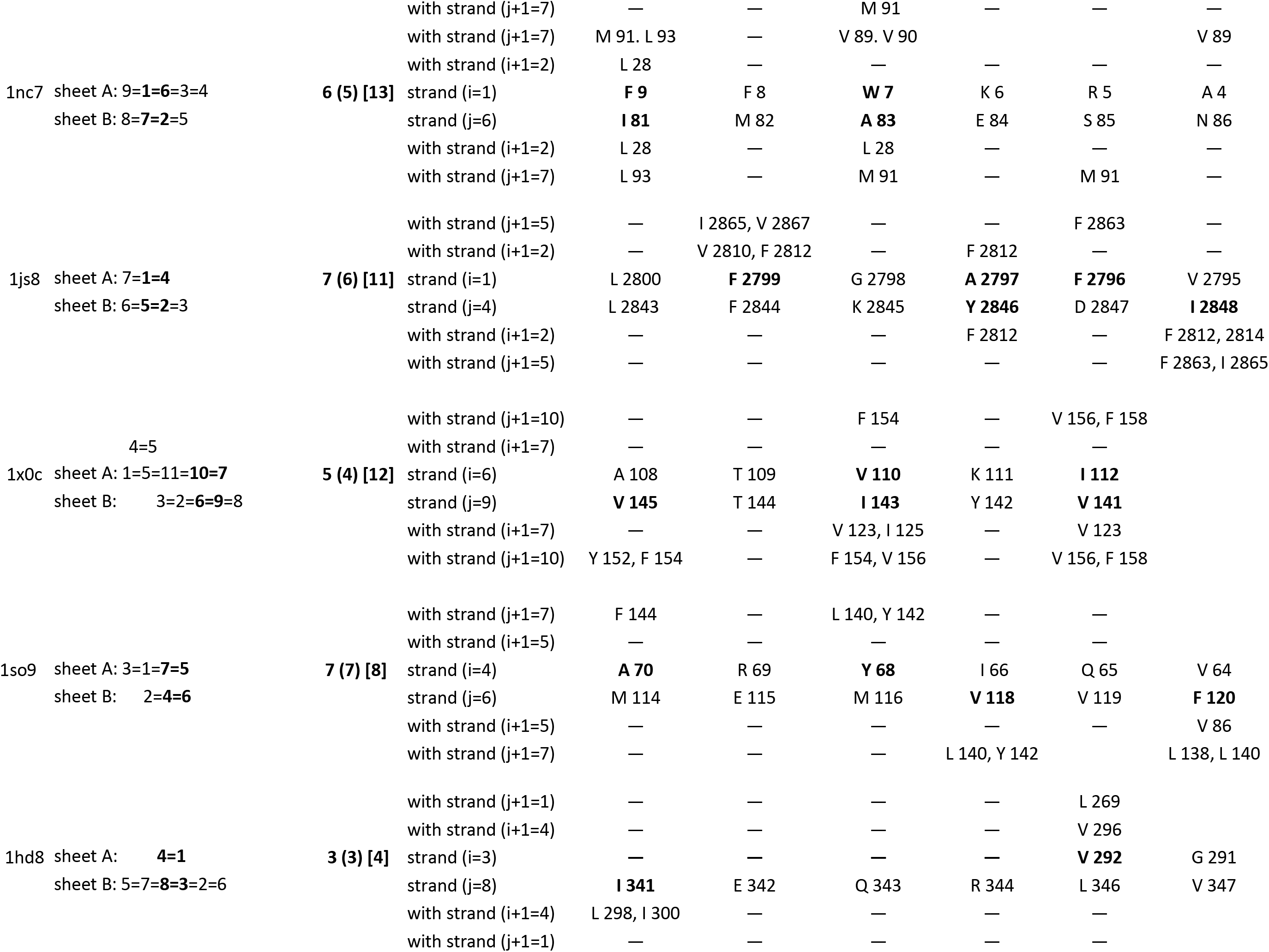
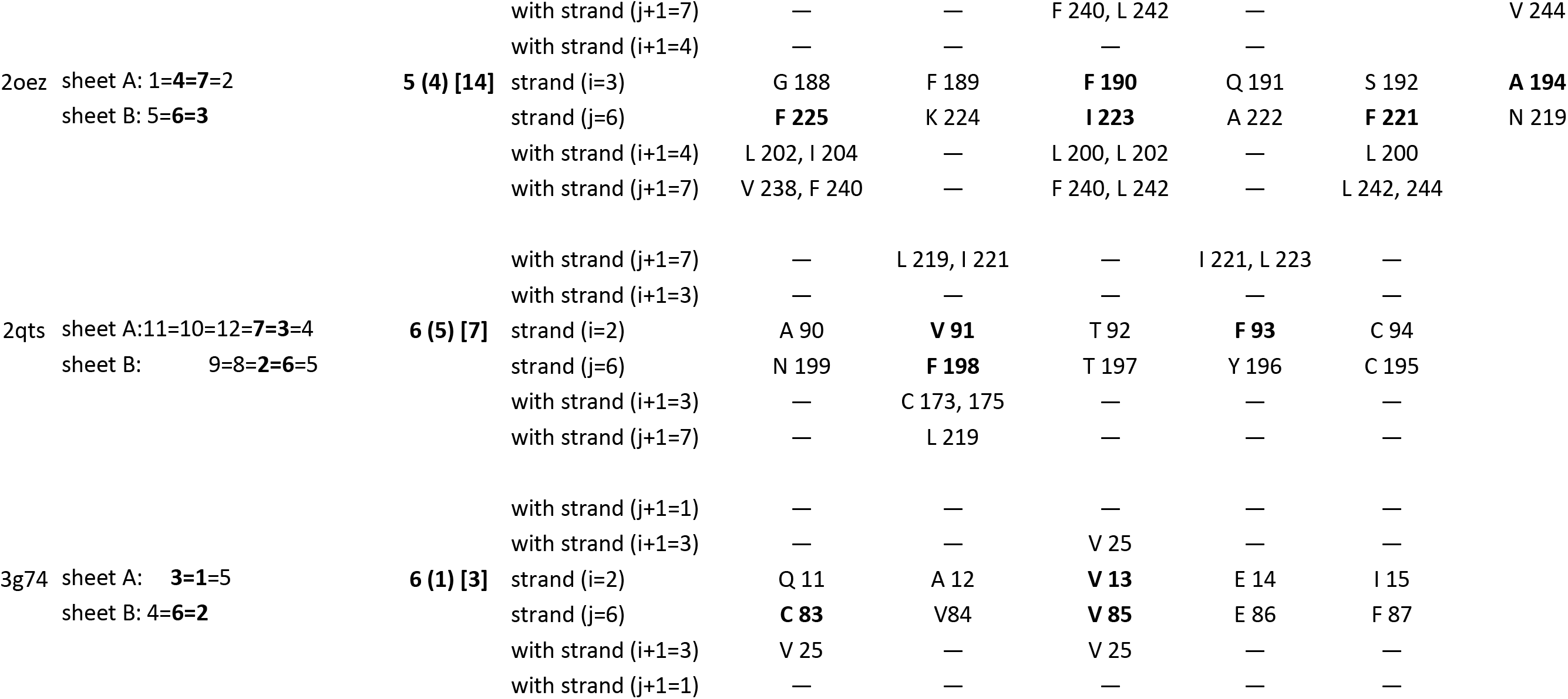
Intersheet contacts of hydrophobic residues in the substructures 2a. The legend is similar to the legend of Table 3a

**Table 4b.** Intersheet contacts of hydrophobic residues in the substructures 2. The legend is similar to the legend of Table

| PDB code | SubS 2b | Contacts in SubS | Intersheet Contacts | Residue- residue intersheet contacts (PHR-IC) |  |  |  |  |  |  |
| --- | --- | --- | --- | --- | --- | --- | --- | --- | --- | --- |
| 1dlm | sheet A: **1=5=4=7=8=9**<br>sheet B: **2=6=3=11=10** | **2 (2) [7]** | with strand (j+1=6) | — | — | — | — | — | — | — |
|  |  |  | with strand (i+1=2) | Y 120 | — | — | — | — | — | — |
|  |  |  | strand (i=1) | **V 118** | S 117 | E 116 | P 115 | A 114 | — | — |
|  |  |  | strand (j=5) | I 183 | **I 182** | S 181 | R 180 | R 179 | **M 178** | — |
|  |  |  | with strand (i+1=2) | — | 120, A 12 | — | — | — | M 123 | — |
|  |  |  | with strand (j+1=6) | — | 190, V 19 | — | — | — | I 195 | — |
| 3tj8 | sheet A: **1=5=4**<br>sheet B: **2=6=7=3** | **4 (3) [7]** | with strand (j+1=6) | — | — | L 63, F 64 | — | F 64 | — | — |
|  |  |  | with strand (i+1=2) | — | — | — | — | — | — | — |
|  |  |  | strand (i=1) | R 5 | E 4 | **I 3** | I 2 | **M 1** | — | — |
|  |  |  | strand (j=5) | **L 49** | R 48 | **I 47** | A 46 | **I 45** | — | — |
|  |  |  | with strand (i+1=2) | — | — | — | — | — | — | — |
|  |  |  | with strand (j+1=6) | L 63 | — | L 63 | — | L 63, F 64 | — | — |
| 1x0c | 4=5<br>sheet A: **1=5=11=10=7**<br>sheet B: **3=2=6=9=8** | **4 (3) [7]** | with strand (j+1=6) | — | A 108 | — | — | — | — | — |
|  |  |  | with strand (i+1=2) | — | V 59 | — | — | — | — | — |
|  |  |  | strand (i=1) | Q 27 | **L 28** | L 29 | T 30 | W 31 | — | — |
|  |  |  | strand (j=5) | S 105 | **Y 104** | L 103 | **F 102** | S 101 | T 100 | W 99 |
|  |  |  | with strand (i+1=2) | — | — | — | V 57, V 59 | — | — | — |
|  |  |  | with strand (j+1=6) | — | A 108 | — | 108, V 11 | — | — | — |

**Table 4c.** Intersheet contacts of hydrophobic residues in the substructures 2c. The legend is similar to the legend of Table 3a

| PDB code | SubS 2c | Contacts in SubS | Intersheet Contacts | Residue- residue intersheet contacts (PHR-IC) |  |  |  |  |  |
| --- | --- | --- | --- | --- | --- | --- | --- | --- | --- |
| 1h8e | sheet A: 2=1=9=8=5<br>sheet B: 4=3=6=7=10 | 4 (1) [2] | with strand (j=9) | A 94 | — | — | — | — | — |
|  |  |  | with strand (i+1=8) | — | — | — | — | — | — |
|  |  |  | strand (i=7) | V 77 | F 76 | Y 75 | K 74 | S 73 | T 72 |
|  |  |  | strand (j+1=10) | A 97 | V 98 | T 99 | L 100 | — | — |
|  |  |  | with strand (i+1=8) | — | — | — | — | — | — |
|  |  |  | with strand (j=9) | A94 | — | — | — | — | — |
| 1g6e | sheet A: 1=3=2=5=8<br>sheet B: 4=9=6=7 | 1 (1) [6] | with strand (j=9) | — | I 86 | I 84, I 86 | — | — | — |
|  |  |  | with strand (i+1=4) | — | — | — | — | — | — |
|  |  |  | strand (i=3) | N 28 | A 27 | F 26 | C 25 | L 24 | — |
|  |  |  | strand (j+1=1) | I 2 | N 3 | R 4 | T 5 | — | — |
|  |  |  | with strand (i+1=4) | M 32 | — | — | — | — | — |
|  |  |  | with strand (j=9) | I 84, I 86 | — | — | — | — | — |
| 1knb | sheet A: 10=3=2=1=5=6<br>sheet B: 4=9=8=7 | 6 (6) [5] | with strand (j=9) | W 556 | — | W 556 | — | — | — |
|  |  |  | with strand (i+1=4) | — | — | — | — | — | — |
|  |  |  | strand (i=3) | V 438 | S 437 | V 436 | T 435 | A 434 | I 432 |
|  |  |  | strand (j+1=10) | — | — | V 573 | T 574 | F 575 | Y 577 |
|  |  |  | with strand (i+1=4) | — | — | — | — | — | — |
|  |  |  | with strand (j=9) | — | — | — | — | M 552, F 554 | M 552 |
| 2apn | sheet A: 1=6=7<br>sheet B: 3=2=5=4 | 4 (1) [2] | with strand (j=5) | — | L 69 | — | — | — | — |
|  |  |  | with strand (i+1=2) | — | — | — | — | — | — |
|  |  |  | strand (i=1) | T 10 | F 11 | T 12 | D 13 | A 14 | — |
|  |  |  | strand (j+1=6) | G 82 | T 83 | V 84 | D 85 | Y 86 | — |
|  |  |  | with strand (i+1=2) | — | — | — | — | — | — |
|  |  |  | with strand (j=5) | — | — | L 69 | — | — | — |
| 1n7v | sheet A: **1=6=7=8=9**<br>sheet B: **5=2=3=4** | **3 (2) [7]** | with strand (j=5) | — | — | — | — | — | — |
|  |  |  | with strand (i+1=2) | — | — | — | A 51, L 52 | — | — |
|  |  |  | strand (i=1) | — | — | N 43 | **Y 44** | W 45 | G 46 |
|  |  |  | strand (j+1=6) | **Y 355** | **A 356** | V 357 | **Y 358** | K 359 | T 360 |
|  |  |  | with strand (i+1=2) | — | A 54 | — | L 52, A 54 | — | — |
|  |  |  | with strand (j=5) | A 353 | A 353 | — | — | — | — |

**Table 4d.**
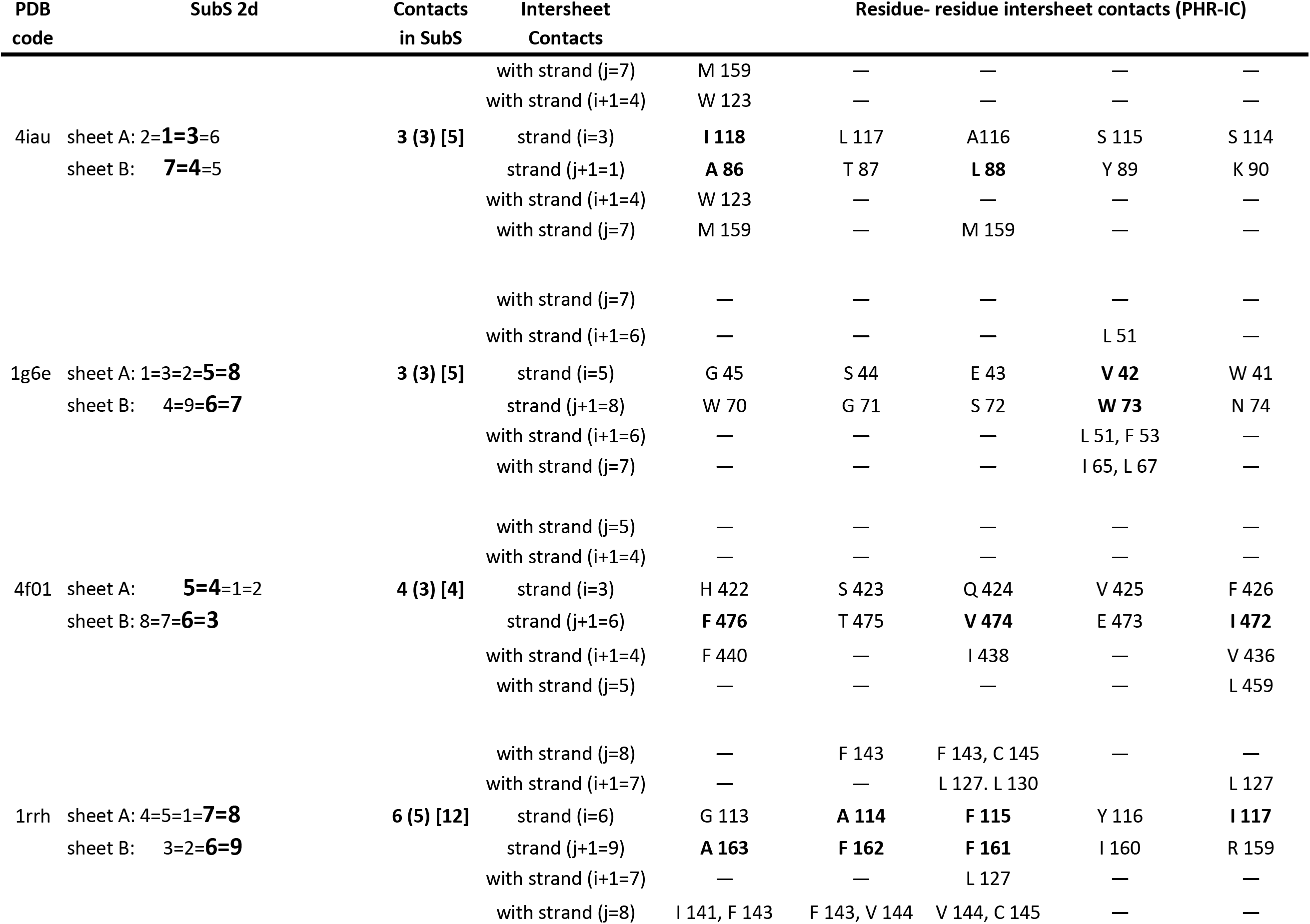
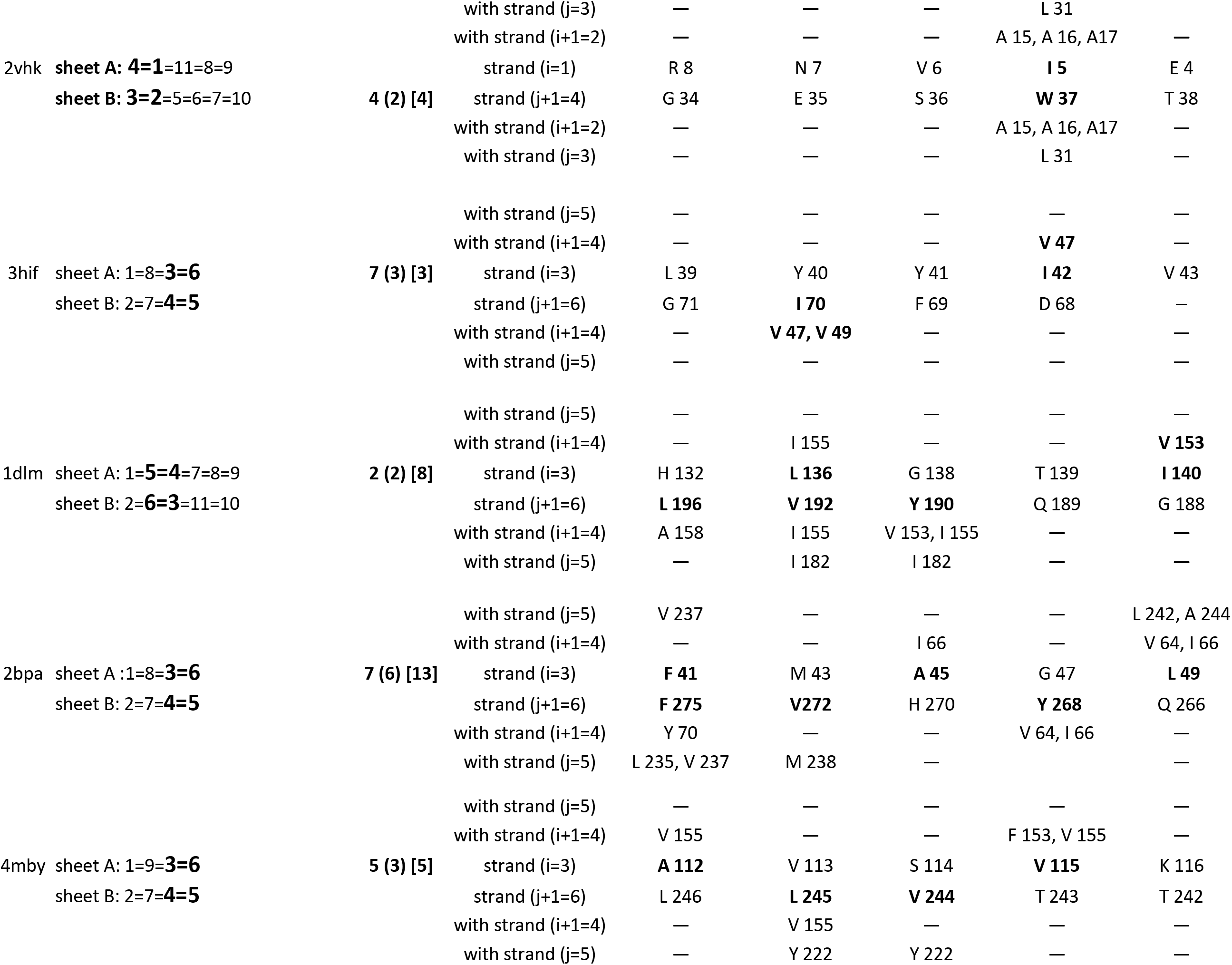
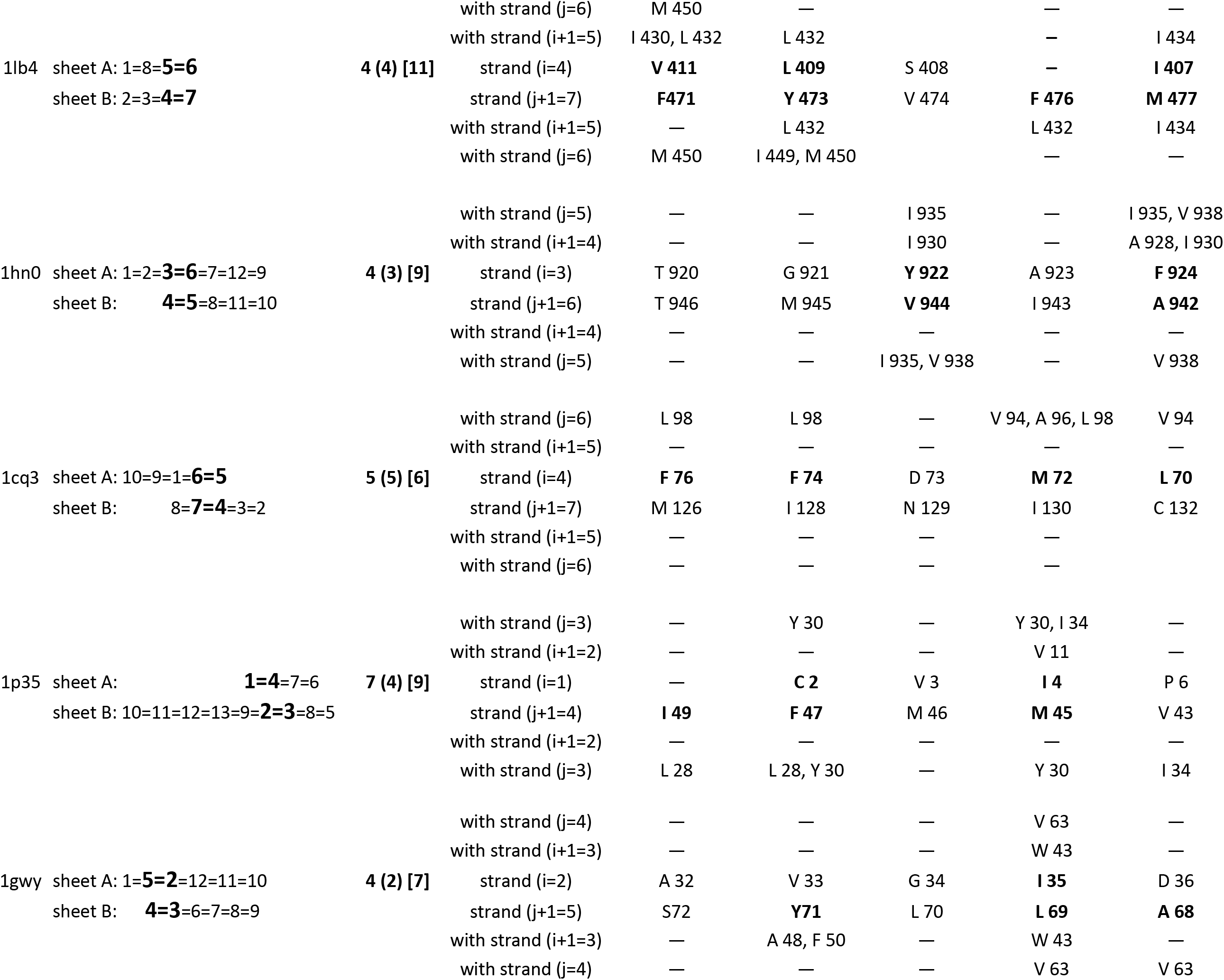
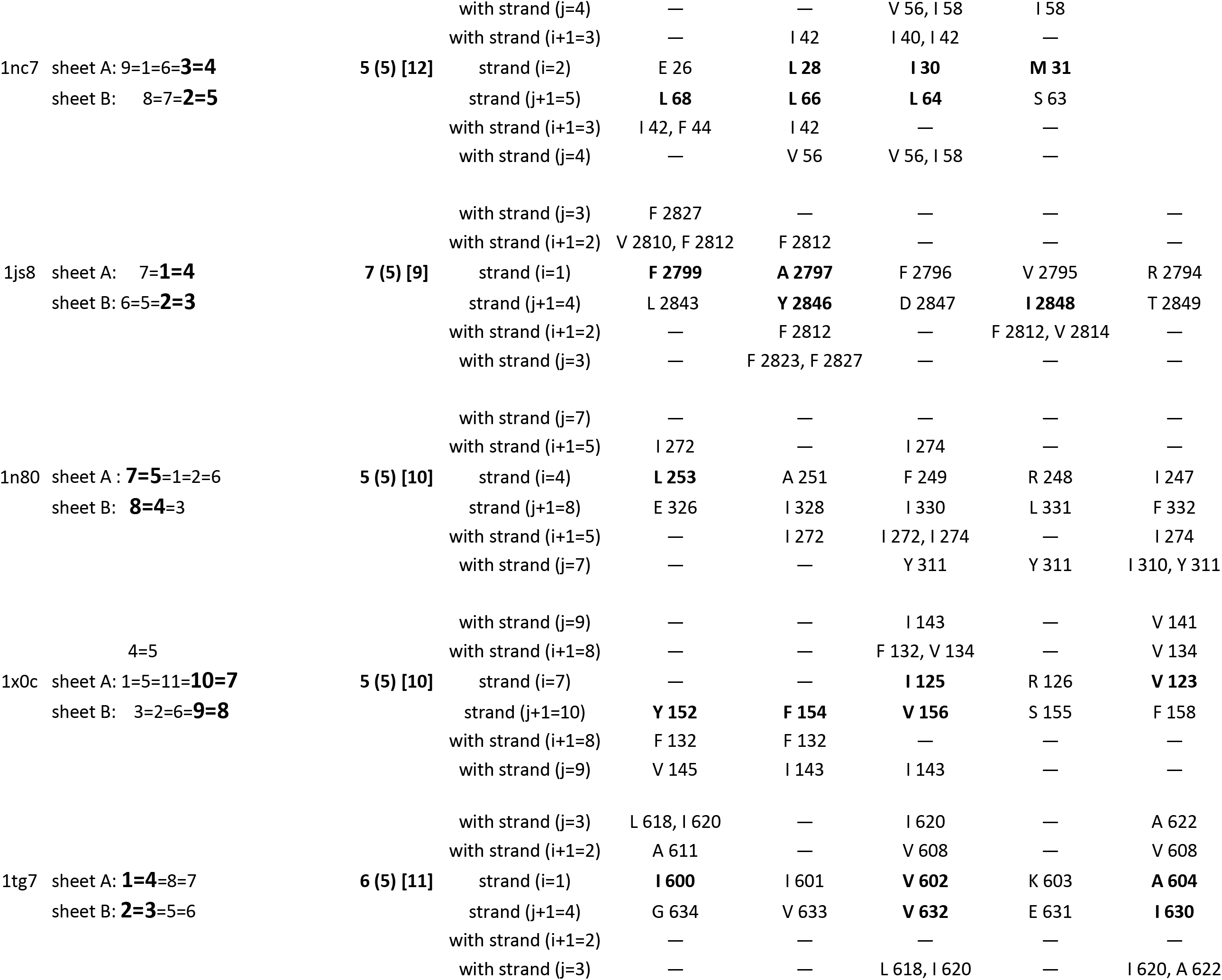
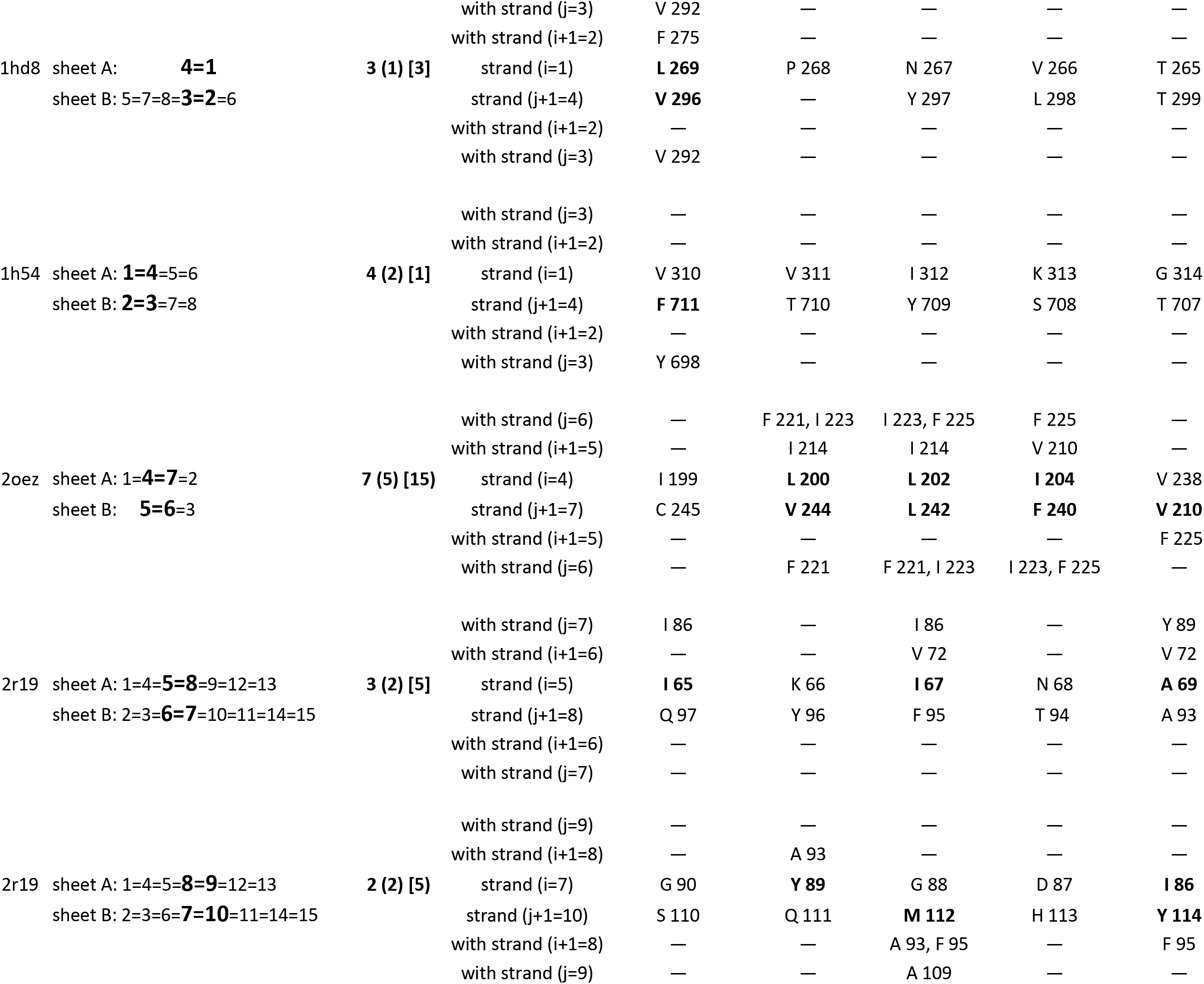
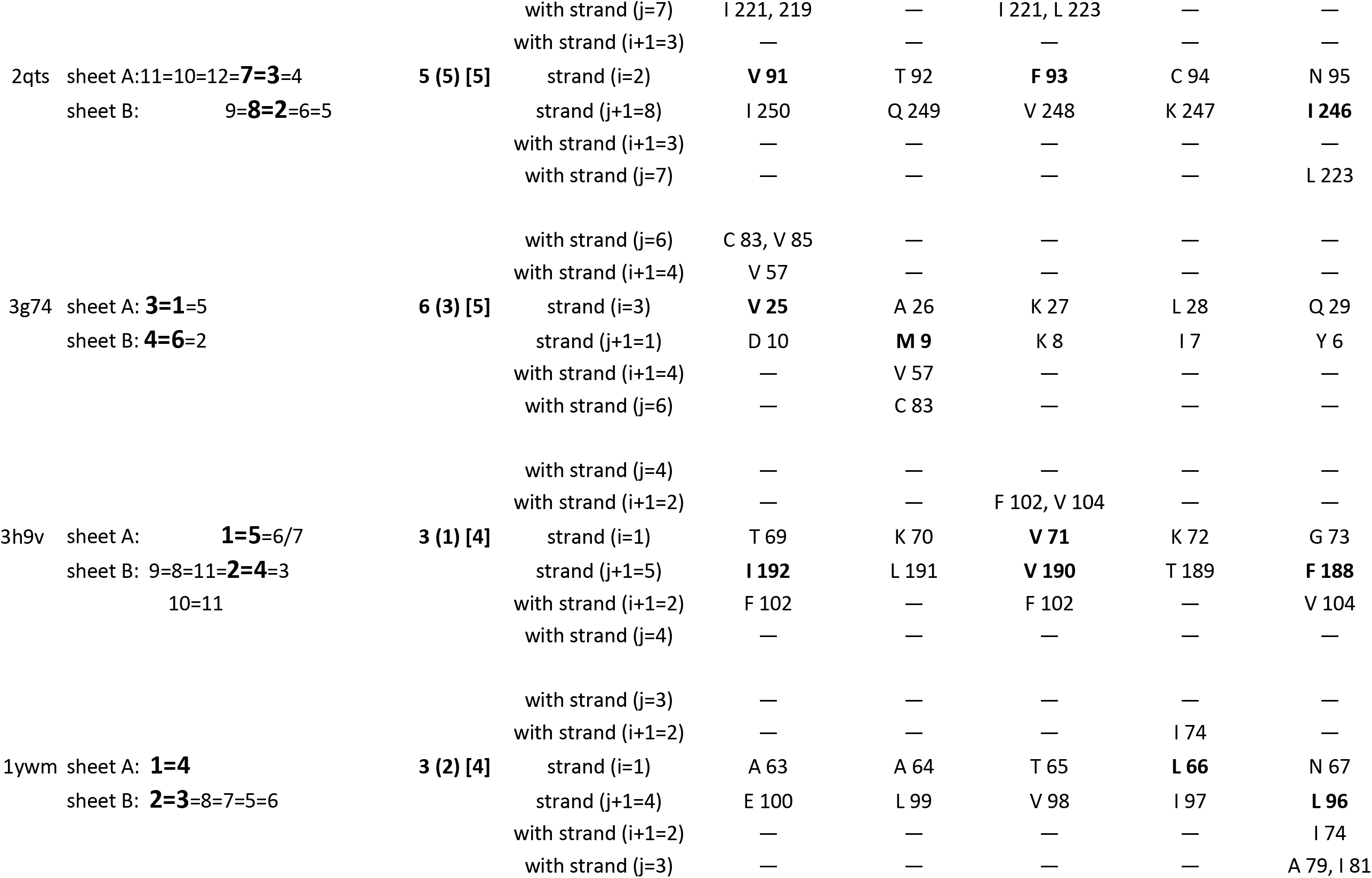
Intersheet contacts of hydrophobic residues in the substructures 2d. The legend is similar to the legend of Table 3a

## Discussion

### Prerequisites for the formation of sandwich structures

All analyzed structures with the sandwich architecture contained at least one conserved substructure of the type shown in Fig. 1 and 2. Thus, the two types of substructures, with their specified arrangements of four strands, may be considered as a prerequisite for the formation of two sheets in sandwich-like protein domains. It is plausible to suggest that the strands are reorganized in space to form the substructure(s) as the critical early step in the formation of hydrophobic core between sheets. The sandwich substructures can be considered as the structural nucleus of protein domains, while non-substructure strands are the ‘annexes’ of protein substructure(s). The presence of the structural nucleus provides indirect evidence that proteins of various topologies in sandwich architecture are either evolutionarily related or are the result of convergent evolution, in which two types of substructure independently formed as indispensable structural elements of stable sandwich domains.

### Similar sequence characteristics of substructures

Identification of the common supersecondary structure characteristics of sandwich proteins - the substructures - allowed for the determination of the common sequence characteristics of sandwich proteins. These common sequence characteristics are the specific distribution of hydrophobic residues in sequences of proteins with sandwich architecture. The essential role of hydrophobic residue interactions in determining the pathway of protein folding is supported by many studies.^12, 19^

The distribution of hydrophobic residues in the substructure strands in the same beta-sheet can be characterized with respect to:

a. The proportion of hydrophobic residues with a tendency to form beta strands (‘preferred’ residues). The ratios of preferred and non-preferred residues for strands, adjusted for strand length of the strands were considered in our recent work20;
b. A tendency to form PHR and pairs of oppositely charged residues. Ability for form pairs of hydrophobic or charged residue contacts depends on relative positions of respective residues in the neighboring strands in a beta sheet. This imposes great restrictions on the distribution of these residues in the sequence.
c. The proportion of hydrophobic residues located in ‘inward-oriented’ positions in beta-strands. Most residues involved in PHR were found in these inward-oriented positions and formed PHR-IC.

It is can be argued that the similarity of sequences in sandwich substructures is determined not only by the number of conserved positions with similar residues as a result of artificial alignment procedures, but rather by the distributions of hydrophobic residues in the substructure strands in conformance with the above requirements. The algorithm for the distribution of residues in substructure sequences also impose certain limitations on the arrangement of hydrophobic residues in the non-substructure strands (see framed PHRs in the columns ‘sequences of strands in beta-sheets A and B” in Table 2a, b).

The specific rules of distribution of hydrophobic residues within the substructures are characteristics of sandwich proteins but are not a sufficient criterion to classify structure as having sandwich architecture. The sequence template for a particular sandwich topology includes not only the rules of distribution of hydrophobic residues within the substructure strands but also the specifications of the minimum required the number of residues with high-frequency adjusted for the lengths of strands and loops respectively. These data can be obtained from statistical analysis of residues of strands and loops in all beta proteins. It was shown that these sequence characteristics are very specific and sensitive for identifying proteins of a given fold from all known protein sequences.^20^

#### Beta-Protein linguistics in substructures

The term ‘protein linguistics’ is often used due to the numerous parallels between linguistics and biology.^21–29^ Human linguistics analyzes how letters of the alphabet form meaningful texts, while protein linguistics analyzes how a sequence made up of 20 amino acids ‘letters’ forms a stable and biologically active three-dimensional structure. In linguistics, as in biology, the essential problem is to identify laws that determine the nonrandom arrangement of elements in sequences. Linguistics usually considers a three-level scheme of analysis: Letter ◊ Word ◊ Sentence. These levels of analysis can be applied to biological languages as well. The letters are conditional symbols that designate sounds in written human languages or the monomers of polypeptide chains in protein languages.

The problem of defining ‘word’ and ‘sentence’ is more complex. One of the best-known definitions for a word in linguistics was introduced by Bloomfield: the word is the “smallest free form”, i.e. a form that can occur alone as an expression.^36^ In the protein linguistics, the definitions of ‘word’ and ‘sentence’ differ depending on the objectives of the study. For example, to analyze the location of domains in a multi-domain structure, domains can be considered ‘words’, and the tertiary structure of a protein - ‘sentence’.^37–38^ If the goal is to analyze the sandwich-like domain then the protein domain can be regarded as a ‘sentence’.

For our purposes, in beta-proteins linguistics two adjacent strands in a beta-sheet may be considered a ‘word’ because they make up the smallest part of the beta-sheet that can exist independently. Thus, in beta-protein morphology, the ‘word’ consists of two sequence fragments that are separated by a string of residues. (Note, in alpha protein linguistics a single helix can be considered as a ‘word’.)

Our study focused on a four-strand “mini-sandwich” substructure that can exist as an autonomous domain or as part of a domain with more than four strands. Such a substructure can be regarded as an independent ‘clause’ in beta-protein linguistics since independent ‘clause’ in human linguistics is a part of the sentence that can stand alone as a sentence.

The next level of language complexity is syntax ─ rules that govern the arrangement of words to create grammatically correct clauses and sentences. Protein syntax determines the formation of a substructure and, ultimately, a stable and functionally viable three-dimensional structure. If a word consists, for example, of two strands *i* and *j* (Fig. 2a) then а ‘half-word’ - strand *i* - may be a neighbor of the other strand *m* which participates in the formation of another word: *m* – *i*. Thus, one strand could be a part of two words, e.g. strand *j* can be part of *i* – *j* word as well as *j* – *k* word. Therefore, the beta-sheet is a set of words that are partially enclosed in each other. This design fortifies the structure of the beta-sheet.

We can deduce some of the rules that govern the process of converting a 1D sequence of residues into ‘words’ and then into a ‘meaningful’ 3D sandwich structure that is stable and capable of performing biological functions. These rules can be formulated as follows:

1. For each beta-protein word (a pair of H-bonded strands in a beta-sheet) one can estimate the minimum allowed the number of letters (residues) favorable for word formation and the maximum allowed the number of letters that are unfavorable to word formation. These conditions are determined based on the statistical analyses of residues in strands in all beta proteins.
2. Whether a sequence fragment can form a word will depend on the coordinated arrangement of hydrophobic residues in a fragment, which will determine the possibility of PHR formation in a word. The coordinated arrangement of pairs of oppositely charged residues is also frequently observed in words.
3. Syntax rules determine the relative position of words in two beta sheets to ensure the stability of the substructure. The condition is the location of most hydrophobic residues in both words at positions in which the residues can form intersheet contacts. In all examined substructures there is at least one PHR in which both residues are directed inward.

## Material and Methods

### Structural analysis

The sandwich architecture in the CATH database encompasses 44 protein folds^1^. Proteins of two folds (2.60.460 and 2.60.530) were not included because structural classification in PDBSum database^30^ does not support sandwich-like architecture for proteins in these two folds. Thus, 42 folds, which describe 528 protein superfamilies, about 40% of all beta-protein superfamilies, were interrogated. One protein domain from every fold, which served as the representative for its respective fold in the CATH database, was examined. For secondary structure analysis, the information about localization of strands in sequences was taken from PDB.^31^ For supersecondary structure (SSS) of the domains, PDBsum topology diagrams that show the arrangement and connectivity of strands in beta sheets were used.

The strands in the domains were numbered starting at the N-terminus. The SSSs are presented according to their arrangements in the two beta sheets (Table 1, column ‘SSS domain’). For residue-residue contact analysis, CSU software was used, which calculates contact surface area for all atoms in residues and provides detailed description of contacts.^32^

#### Definition of hydrophobic residues

Our analysis focuses on hydrophobic residues Ile, Val, Leu, Phe, Met, and Trp in strands. Residues Ala, Tyr, and Cys were added to the group of hydrophobic residues because they often substitute for hydrophobic residues in strands. The weakly hydrophobic Ala is considered both “interior-seeking” and “water-seeking” according to many hydrophobic scales.^33^ Although Tyr has both polar and nonpolar groups, it often substitutes for hydrophobic residues in strands and is classified as a hydrophobic amino acid on most hydrophobicity scales.^34^ Cys residue can play the role of a hydrophobic residue in specific protein microenvironments.^35^

